# The Structural Role of N170 in Substrate-assisted Deacylation in KPC-2 β-Lactamase

**DOI:** 10.1101/2023.05.22.541848

**Authors:** Diksha Parwana, Jing Gu, Shuang Chen, Christopher R. Bethel, Emma Marshall, Andrea M. Hujer, Robert A. Bonomo, Shozeb Haider

**Affiliations:** UCL School of Pharmacy, London, UK; Research Service, Louis Stokes Cleveland Department of Veterans Affairs Medical Center, Cleveland, OH, USA; Department of Medicine, Case Western Reserve University School of Medicine, Cleveland, OH, USA; Clinician Scientist Investigator, Louis Stokes Cleveland Department of Veterans Affairs Medical Center, Cleveland, OH, USA; Department of Molecular Biology and Microbiology, Pharmacology, Biochemistry, and Proteomics and Bioinformatics, Case Western Reserve University School of Medicine, Cleveland, OH, USA; and CWRU-Cleveland VAMC Center for Antimicrobial Resistance and Epidemiology (Case VA CARES) Cleveland, OH, USA; UCL Centre for Advanced Research Computing, London, UK

**Keywords:** Ceftazidime, Moxalactam, β-lactamase, metadynamics, substrate-assisted catalysis

## Abstract

The amino acid substitutions in *Klebsiella pneumoniae* carbapenemase 2 (KPC-2) that have arisen in the clinic are observed to lead to the development of resistance to ceftazidime-avibactam, a preferred treatment for KPC bearing Gram-negative bacteria. Specific substitutions in the omega loop (R164-D179) results in changes in the structure and function of the enzyme, leading to alterations in substrate specificity, decreased stability, and more recently observed, increased resistance to ceftazidime/avibactam. Using accelerated rare-event sampling well-tempered metadynamics simulations, we explored in detail the structural role of R164 and D179 variants that are described to confer ceftazidime/avibactam resistance. The buried conformation of D179 substitutions produce a pronounced structural disorder in the omega loop - more than R164 mutants, where the crystallographic omega loop structure remains mostly intact. Our findings also reveal that the conformation of N170 plays an underappreciated role impacting drug binding and restricting deacylation. The results further support the hypothesis that KPC-2 D179 variants employ substrate-assisted catalysis for ceftazidime hydrolysis, involving the ring amine of the aminothiazole group to promote deacylation and catalytic turnover. Moreover, the shift in the WT conformation of N170 contributes to reduced deacylation and altered spectrum of enzymatic activity.

## Introduction

β-lactams, including penicillins, cephalosporins, carbapenems, and monobactams, are the most commonly used antibiotics that irreversibly inhibit enzymes involved in bacterial cell wall synthesis.^[1]^ As a result of persistent exposure to β-lactams and the imposed survival pressure, bacteria continually evolve to evade therapeutic action, predominantly via the production of β-lactamases in Gram-negative bacteria.^[2–5]^ β-Lactamases, categorized into classes A, B, C, and D,^[6]^ can degrade all classes of β-lactam drugs and thus present a global clinical threat in the form of antimicrobial resistance (AMR).^[7,8]^ To combat the ever-increasing AMR, the use of β-lactamase inhibitors-clavulanic acid, tazobactam, and sulbactam that share the β-lactam backbone, were clinically approved in combination with specific β-lactam drugs.^[9]^ The emergence of *Klebsiella pneumoniae* carbapenemase (KPC) producing strains, prompted the development of novel β-lactamase inhibitors.^[10]^ Efforts to restore activity against these diverse enzymes led to the clinical approval of a non-β-lactam (diazabicyclooctane) β-lactamase inhibitor, avibactam.^[11]^ Unlike the first three clinical inhibitors that are limited in their activity only to class A β-lactamases, avibactam is a broad-spectrum inhibitor for serine (class A, C, and some class D) enzymes.^[8,12]^

The combination of avibactam with an oxyimino-cephalosporin, ceftazidime, appeared as a salvage therapy for the treatment of carbapenem-resistant Enterobacterales (CRE) infections primarily caused by KPC-producing bacteria.^[13]^ However, a site-directed mutagenesis study revealed ceftazidime-avibactam (CAZ-AVI) resistance in KPC-2 variants with substitutions at R164 and D179 positions of the omega loop.^[14,15]^ Similar resistance was reported for an *in vitro* study for KPC-3 expressing Enterobacteriaceae, primarily due to a D179Y substitution.^[16]^ Shortly after, CAZ-AVI resistance was demonstrated in clinical isolates being treated for KPC-expressing CRE infections with this β-lactam-inhibitor combination.^[17]^ A series of studies using site-saturation and site-directed mutagenesis uncovered the complex array of substitutions conferring ceftazidime/avibactam resistance.^[18–20]^

The omega loop (residues R164-D179 in KPC-2) (Fig. 1) plays a vital role in the β-lactamase enzymatic mechanism. Following the nucleophilic attack by S70, the acyl-enzyme complex is deacylated by a conserved water molecule that is strategically held in position by omega loop residues E166 and N170. E166 acts as the general base that activates the water molecule for the subsequent attack on the covalent complex.^[21,22]^ However, in light of the development of ceftazidime-avibactam resistance in clinical isolates and the impacted kinetics of ceftazidime deacylation reported by recent experiments, we aimed to uncover the underlying mechanisms of altered enzymatic activity for KPC-2 mutants. This study investigates the perturbed structural dynamics involving the omega loop region as a result of six significant KPC-2 omega loop substitutions-R164S,^[23]^ R164H,^[24–26]^ D179N,^[14,27–30]^ D179Y,^[18,19,23,28,29,31–35]^ D179A,^[36–38]^ and D179Q^[27]^ that have shown to confer resistance against CAZ-AVI combination. Residues R164 and D179 confer stability to the omega loop by interacting via a salt bridge. Thus, a substitution at either position might cause pronounced structural changes to the middle of the loop, harboring critical catalytic residues.

**Figure 1:**
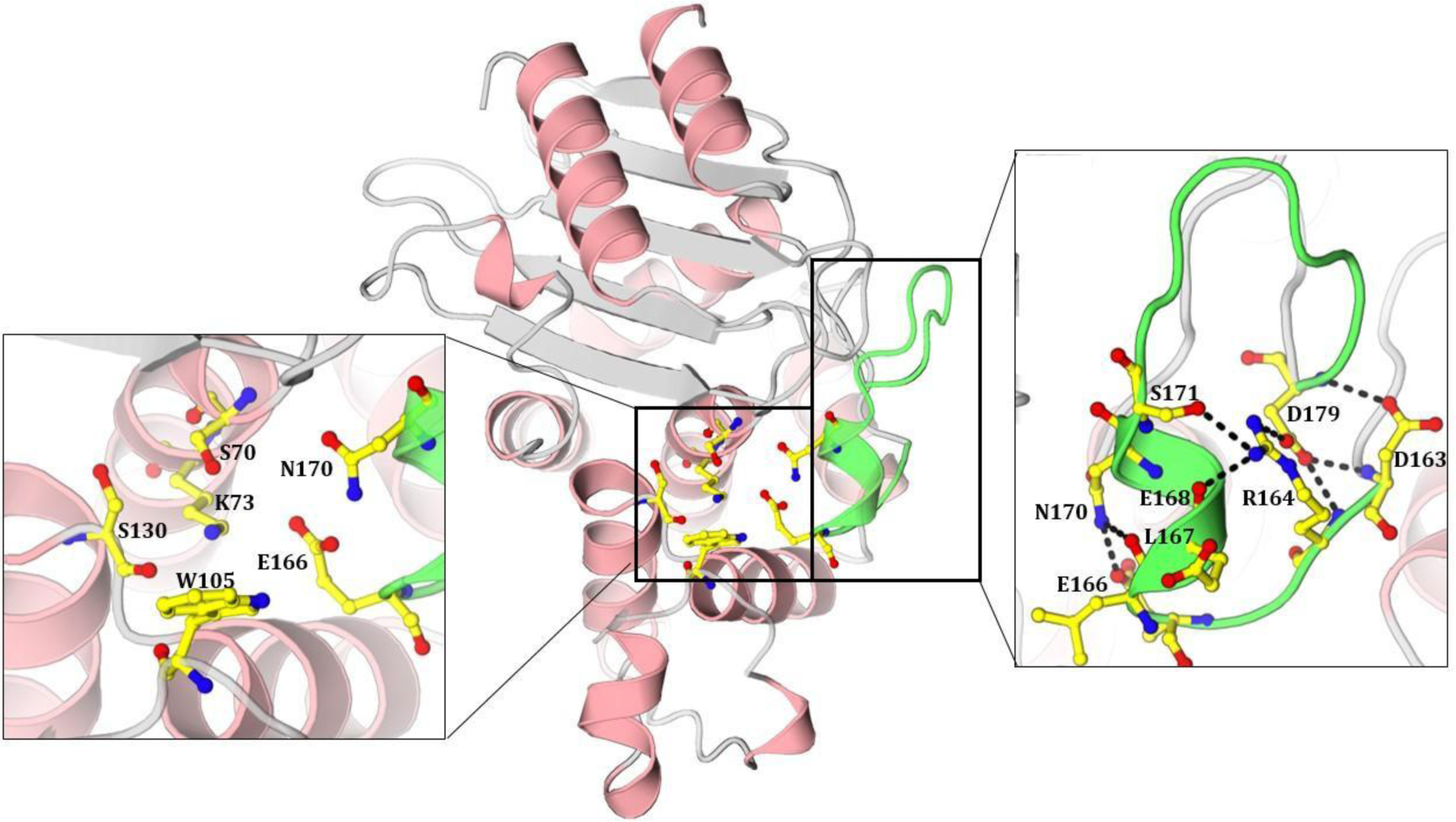
Structure of KPC-2 β-lactamase (PDB id-3DW0) highlighting important active site residues (left) and omega loop interactions (right). The omega loop is colored green.

Using well-tempered metadynamics (wt-MetaD), we accelerated rare-event sampling and uncovered important conformational changes upon R164 or D179 substitution.^[39]^ Due to the buried conformation of D179, variants at this position produce a more pronounced structural disorder in the omega loop compared to R164 mutants, where the crystallographic structure remains mostly intact.^[30,40]^ The findings suggest that the conformation of the omega loop with N170 oriented “away” is favored for KPC-2 omega loop mutants, impacting drug binding and restricting deacylation. This work further strengthens the notion that KPC-2 D179 variants employ substrate-assisted catalysis for ceftazidime, involving the ring amine of the aminothiazole group to promote deacylation and catalytic turnover. Our study attempts to answer the perplexing questions pertaining to the enhanced resistance of KPC-2 mutants to ceftazidime/avibactam intervention as a result of omega loop substitutions. With the emerging resistant strains imposing a major clinical challenge, our results hold significant importance for future drug design.

## Results and Discussion

### D179 variants exhibit altered conformational dynamics

Well-tempered Metadynamics (wt-MetaD) allows us to investigate the most favored conformations of KPC-2 variants, typically separated by large energy barriers and thus challenging to conventional MD simulations.^[39,41,42]^ By forcing the system to escape local minima, wt-MetaD simulations explore the least populated regions of the system within the collective variable (CV) space. The conformations corresponding to the reconstructed free energy minima may shed light on the key regulatory changes and altered enzymatic mechanisms. To understand how mutations alter the dynamics of our system, we characterized the free energy landscape as a function of backbone dihedral angles (φ and ѱ) as they represent the true slow dynamics and could describe global behavior.^[43,44]^ Hence, our wt-MetaD simulations were biased to two CVs-the backbone φ and ѱ dihedral angles of the alerted residue under study, i.e. residue 164 for R164 mutants and residue 179 for D179 mutants. Additionally, two different free energy plots for KPC-2 WT were computed using CVs corresponding to residues R164 and D179.

The free energy surface (FES) for WT as a function of D179-related CVs exhibits one local minimum indicating similar interactions as seen for the native crystal structure (PDB id-3DW0) (Fig 2). Upon back mapping, we observed that the crystal structure lies in the largest free energy minima; thus, it represents the most stable omega loop conformation presenting R164-D179, R164-S171, D179-D163, and E166-N170 interactions.

**Figure 2:**
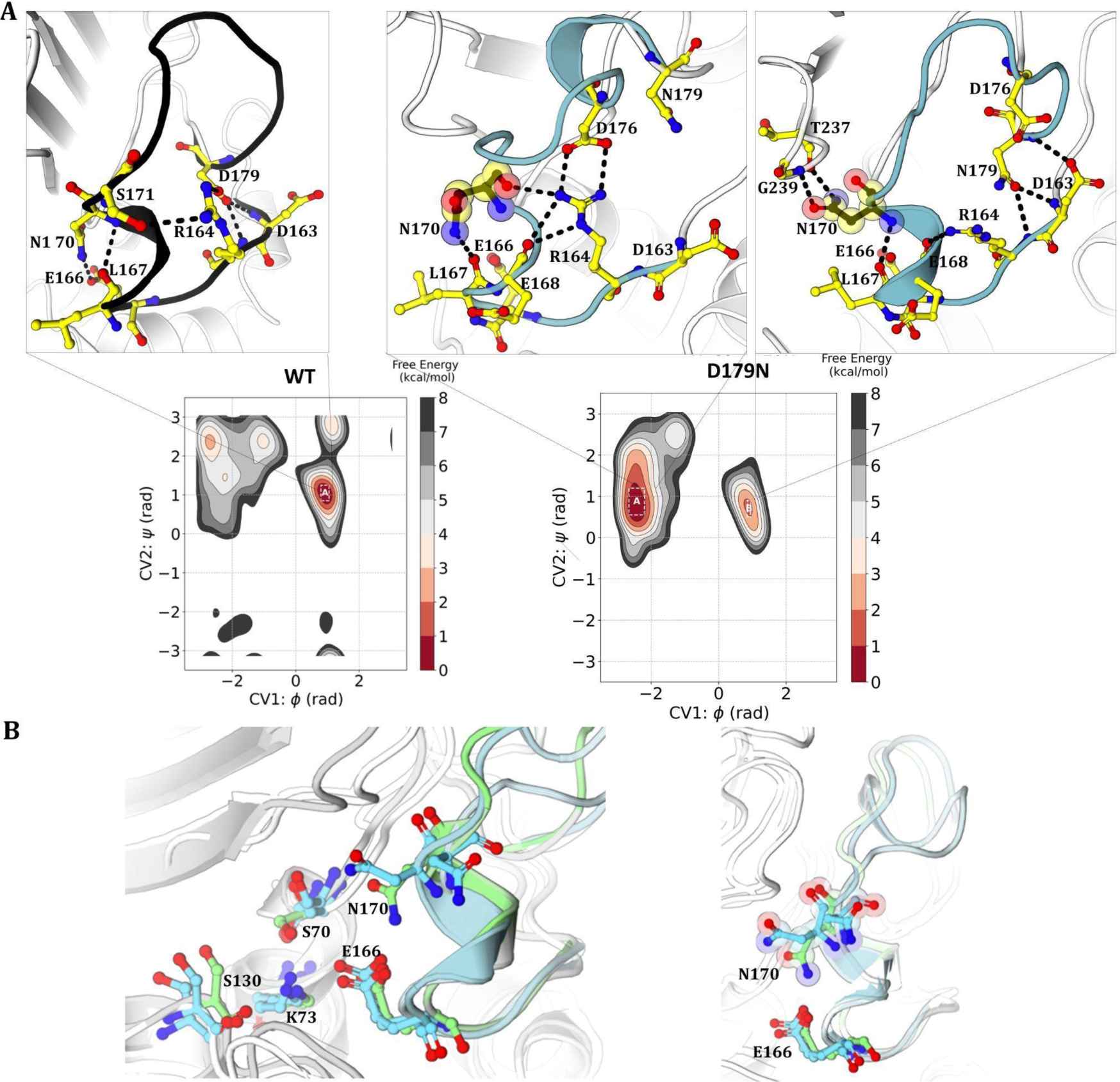
(A) Free energy plots for KPC-2 WT (left) and D179N (right) as a function of φ (CV1) and ѱ (CV2) dihedral angles of residue 179. Omega loop interactions of the conformations extracted from their corresponding free energy minima have been illustrated. (B) Superimposition of two extracted structures from D179N basins with the crystal structure of KPC-2 WT (PDB id-3DW0).

The FES plot for D179N involves two minima as shown in Fig 2. The extracted conformations unravel the disrupted H-bonding network due to the loss of the R164-D179 salt bridge at the neck of the omega loop. In basin B, R164 adopts a solvent-exposed conformation, unable to maintain its native interactions by virtue of D179 substitution and a consequent dynamic loop. Basin A (7 kcal/mol lower in energy than basin B), on the other hand, depicts R164 intact due to the shift in D176 that leads to the formation of the R164-D176 salt bridge. Interestingly, both the conformations sampled by wt-MetaD capture N170 displaced from its crystallographic pose, unable to position the water molecule for effective deacylation of the acyl-enzyme complex. In basin A, the displaced N170 interacts with the backbone of L167, while in the other basin, the shift in the N170 side chain forms new interactions with the backbone of T237 and G238 (consensus numbering). An increased distance from E166 and the formation of new interactions abate the basic character of N170 that is required to activate the deacylating water molecule. Biochemical analysis reporting a decreased rate of deacylation for the hydrolysis of imipenem as well as ceftazidime in D179N strains could be attributed to N170 displacement.^[27]^ The shift in N170 leading to slower hydrolysis further explains the proposition of preferential trapping of β-lactams when compared to avibactam in the expanded active site of D179N mutants.^[27]^ The deacylation mechanism of avibactam in D179N likely involves S130 and K73; thus, its enzymatic kinetics remain unaffected with the shift in N170.^[45]^ Owing to the reduced deacylation rates for β-lactams in the D179N mutant, we suggest the shift in the WT conformation of N170 as the underlying cause for impacted deacylation and altered spectrum of enzymatic activity.

Moreover, N170A substitution in KPC-2 exhibited a > 3000-fold reduction in the deacylation rate for imipenem hydrolysis in a site-directed mutagenesis study.^[46]^ Thus, the crucial involvement of N170 in KPC-2 β-lactamase activity is further supported by the decreased hydrolytic activity upon N170 substitution.

In the D179Y KPC-2 variant, R164 forms a double salt bridge with D176 (basin B and C) or with D163 (basin A) (Fig. 3). Also, in basin A (1 kcal/mol and 1.8 kcal/mol lower in energy than basin B and C respectively), R164-S171 sidechain-sidechain interaction is observed, similar to WT KPC-2. Substitution of the negatively charged carboxyl group with a phenol moiety results in a highly flexible structure owing to the disruption of the H-bond network formed by the buried conformation of D179 in WT. Contrastingly, D179Y adopts an exposed conformation, unable to form interactions within the loop. The lost structural integrity of the omega loop limits the binding of diverse classes of β-lactams in a distorted active site resulting in lower MICs for penicillins (ampicillin, piperacillin), carbapenems (imipenem, meropenem), cephalosporins (cefepime, ceftriaxone, cefotaxime) as well as monobactams (aztreonam).^[19,20]^ The deformed omega loop fails to define the binding pocket anymore and might not be able to sustain the structural changes associated with the binding of a β-lactam in the pocket, thus impeding catalysis. Moreover, the disruption of stabilizing omega loop interactions is unable to constrain N170 in its native conformation involved in the deacylation reaction. Its side chain flips away from the active site and interacts with the L167 main chain (basin B) or remains disengaged (basin A and basin C). The distance of basic carbonyl oxygen of the N170 side chain from the water molecule increases to more than 5 Å in all three observed basins. As a result, slower deacylation might also contribute to the decreased resistance of KPC-2 D179Y mutants to various β-lactams.

**Figure 3:**
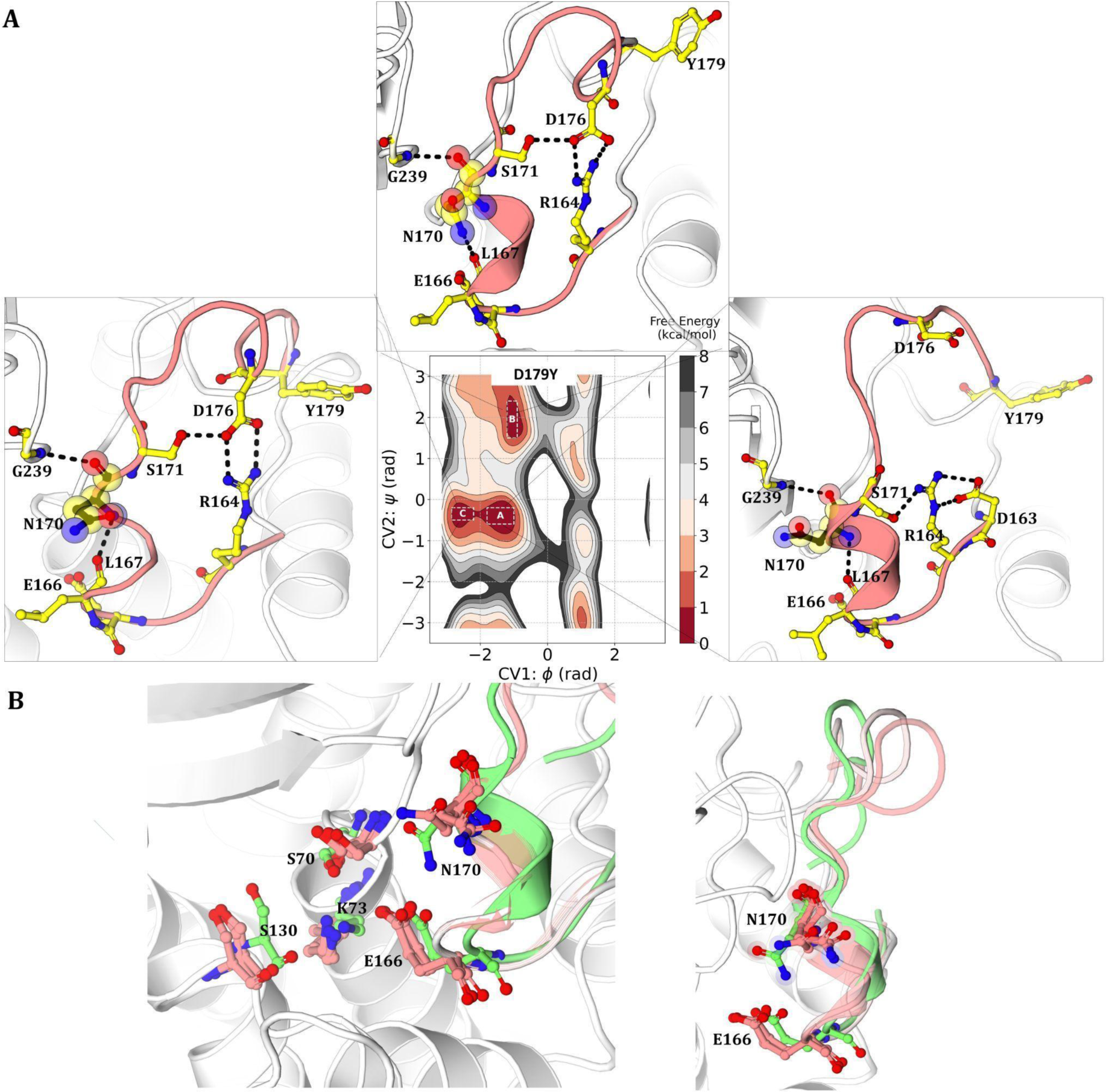
(A) Free energy plot for KPC-2 D179Y as a function of φ (CV1) and ѱ (CV2) dihedral angles of residue 179. Omega loop interactions of the conformations extracted from their corresponding free energy minima have been illustrated. (B) Superimposition of three extracted structures from D179Y basins with the crystal structure of KPC-2 WT (PDB id-3dw0).

In the D179A mutant, as a result of the loss of the R164-D179 salt bridge, the side chain of R178 shifts closer to D163, resulting in a salt bridge as revealed by the most stable conformation, basin A as well as basin C (2.8 kcal/mol higher in energy than basin A) (Fig. 4). In another free energy minima, basin D (3 kcal/mol higher than basin A), R164 shows a new interaction in the form of an ionic bond with the displaced D176 and maintains its native interaction with the sidechain of S171. The helix encompassing E166 and N170 unfolds, and a shift in the side chain of N170 with respect to the WT conformation is observed in all four free energy minima. As a result of the substitution of acidic aspartic acid with hydrophobic alanine, the side chain of N170 orients away from the active site towards the altered residue A179. In the most stable conformation (basin A), the side chain of N170 forms an H-bond with the backbone of A179 and L68. A similar conformation of N170 is observed in basin B as its side chain rotates away from the active site, interacting with the backbone of T180 towards the end of the omega loop. This shift in the N170 side chain, while providing access to an expanded active site, might impede effective deacylation of other β-lactams but not ceftazidime.

**Figure 4:**
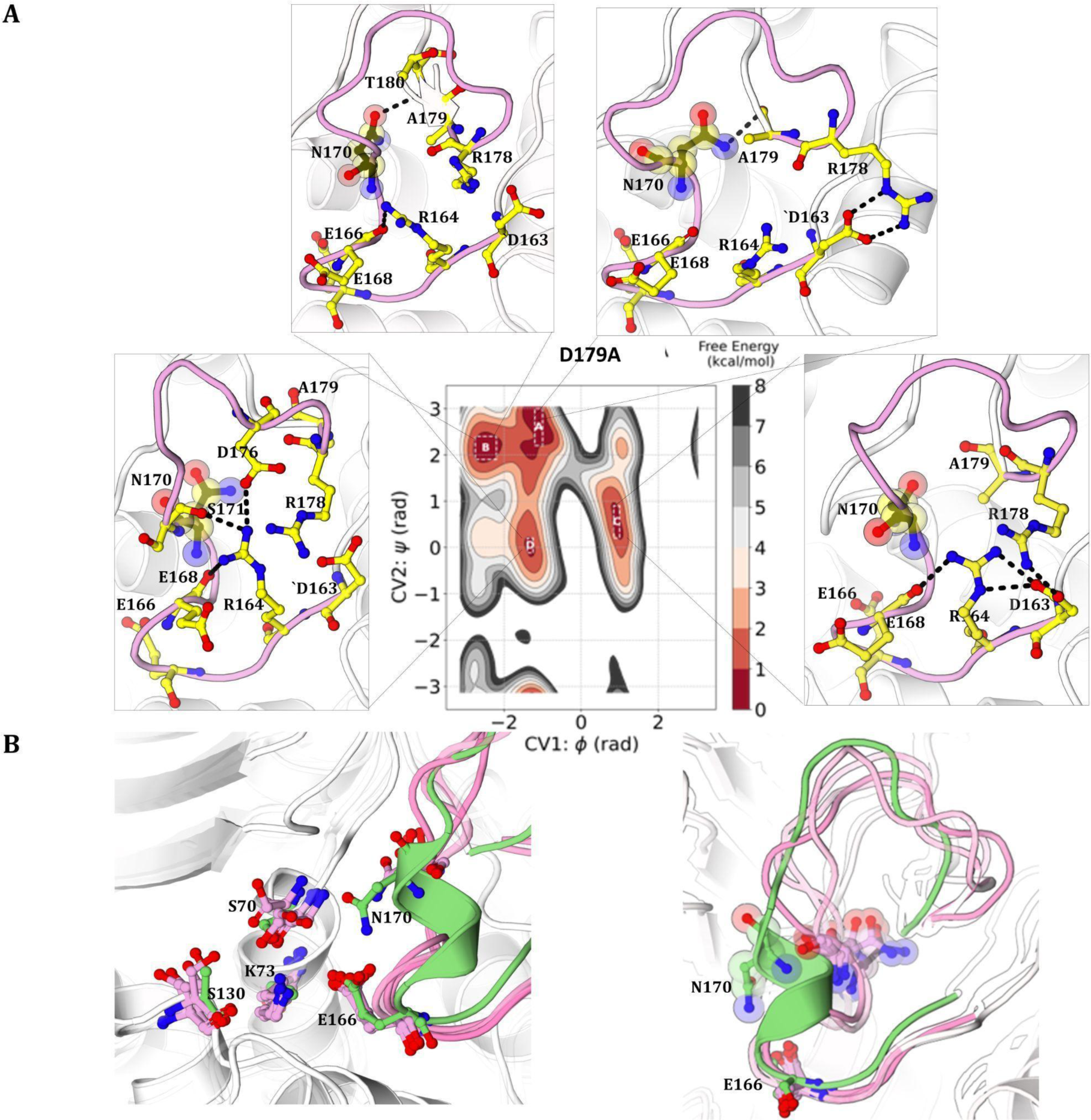
(A) Free energy plot for KPC-2 D179A as a function of φ (CV1) and ѱ (CV2) dihedral angles of residue 179. Omega loop interactions of the conformations extracted from their corresponding free energy minimas have been illustrated. (B) Superimposition of four extracted structures from D179A basins with the crystal structure of KPC-2 WT (PDB id-3dw0).

The substitution of aspartic acid with glutamine in KPC-2 D179Q results in the opening of the omega loop due to the absence of integral bonds at the neck of the structure (Fig. 5). Consequently, R164 fails to stay intact with respect to its WT orientation and either interacts with E168 (basin A and B) or S171 (basin C). The particularly dynamic motions of D176 and Q179 arising due to changes in the conformational equilibrium engender a disintegrated overlying region of the omega loop. Enhanced mobility owing to D179Q mutation in the omega loop is transmitted to the active site residue N170 resulting in the disruption of E166-N170 interaction. Unlike other clinically observed D179 substitutions included in this study, D179Q mutation has not been reported in clinical isolates. Its altered spectrum of activity has been studied using saturation mutagenesis studies *in-vitro*.^[27]^

**Figure 5:**
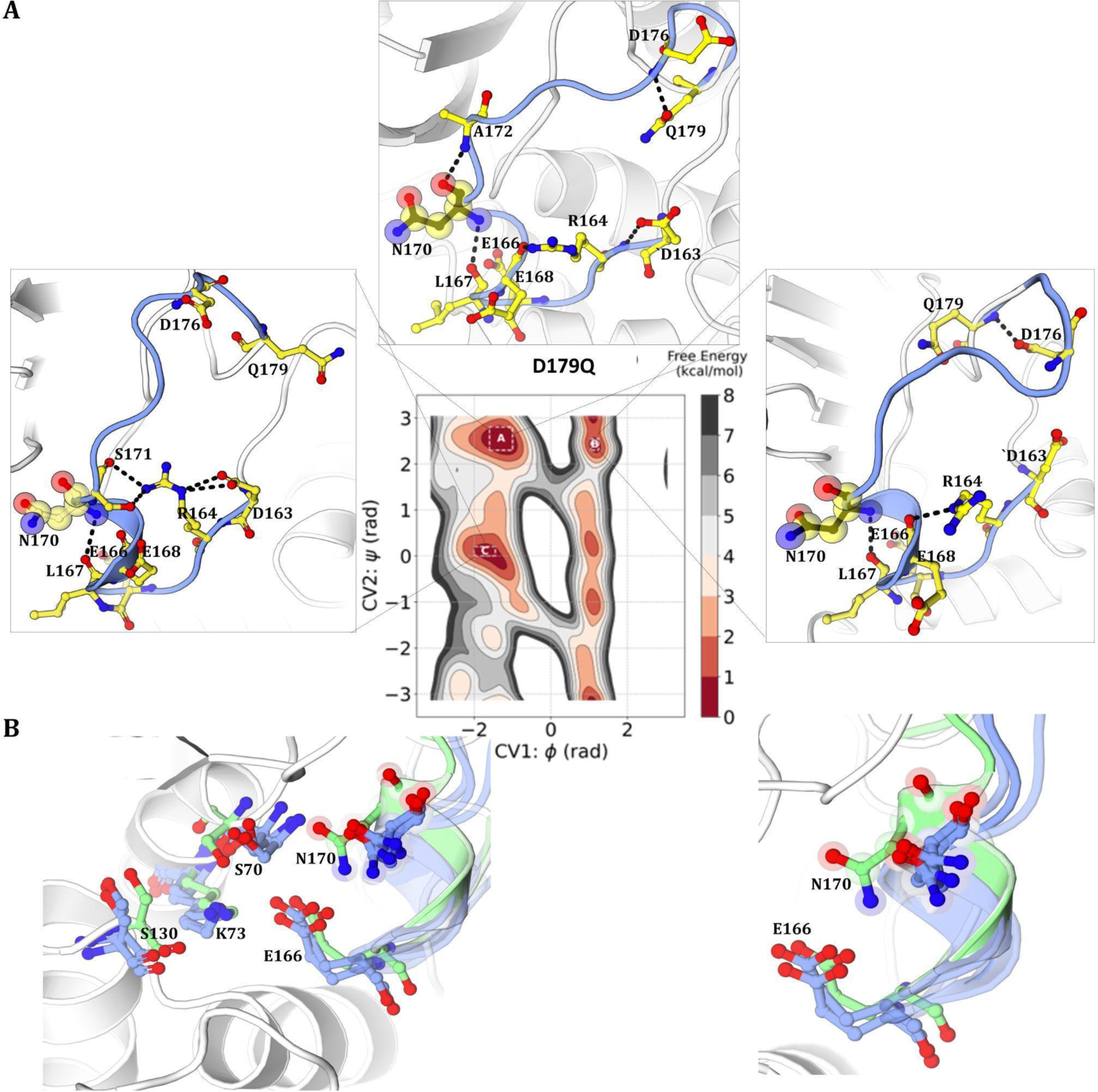
(A) Free energy plot for KPC-2 D179Q as a function of φ (CV1) and ѱ (CV2) dihedral angles of residue 179. Omega loop interactions of the conformations extracted from their corresponding free energy minima have been illustrated. (B) Superimposition of three extracted structures from D179Q basins with the crystal structure of KPC-2 WT (PDB id-3dw0).

For D179 variants (D179N, D179Y, D179A, and D179Q), conformations extracted from the free energy minima outline the enhanced flexibility of the omega loop due to the disruption of the R164-D179 salt bridge. Concomitant with the unstable omega loop is the displacement of the N170 side chain away from the active site, unable to interact with E166 and prime the water molecule for deacylation. While the profound fluctuation of N170 is consistent for each energetically stable conformation for all D179 mutants, the omega loop mutation doesn’t seem to distort E166, which maintains its WT-like conformation facing the active site throughout the sampling time. This leads us to reason the shift in N170 as the underlying cause for lowered deacylation rate of D179 mutants for ceftazidime as well as other β-lactams. As for binding kinetics, the compromised structural integrity of the omega loop renders the active site inefficient to accommodate typically small β-lactam drugs causing D179 variants to become susceptible to several classes of β-lactams. However, increased omega loop flexibility with N170 oriented away is likely to aid in the binding of bulky β-lactam, like ceftazidime and thereby giving rise to enhanced resistance of KPC-2 variants to the ceftazidime-avibactam combination. Results of previous studies on GES type class A β-lactamases reveal the variation in the kinetics of hydrolysis of a non-bulky carbapenem, imipenem, with other comparatively bulkier β-lactams.^[47]^ Our findings reveal the basis for altered substrate specificity for the active site of KPC-2 mutants in the context of rising CAZ-AVI resistance.

### Displaced N170 impacts β-lactamase activity

In the most favored conformations of D179 mutants, the dramatic shift transmitted towards N170, as a result of loss of the R164-D179 salt bridge, is revealed that is deacylation incompetent as well as results in an expanded active site. KPC-2 D179 mutants also present a modest shift in the side chain of S70 by 1.5 Å, whereas other catalytic residues i.e, K73, S130, and E166, maintain their overall WT-like conformations. While D179 mutants develop increased resistance to ceftazidime, their activity against other β-lactam drugs diminishes. Kinetic studies report lower MICs and decreased turnover (*k*_cat_) in the hydrolysis of several penicillins, carbapenems, cephalosporins, as well as monobactams, especially for the D179Y mutant.^[18,19,24,32,33,48]^ This elevated susceptibility of KPC-2 D179 strains to various β-lactams is attributed to the highly disordered omega loop that limits the effective hydrolysis of β-lactams. The disorientation of N170 results in an expanded binding pocket and impedes the deacylation of the acyl-enzyme complex. The displaced S70 side chain might additionally restrict the acylation of the substrate.^[27]^ Paradoxically, the repositioning of the N170 side chain eases the accommodation of the bulky β-lactam, ceftazidime, due to the concomitant expansion of the binding pocket. Docking of ceftazidime in the active site of KPC-2 variants indicates the orientation of the aminothiazole ring of C7 substituent of ceftazidime towards WT conformation of N170, thus highlighting the potential bypass of steric clashes upon N170 displacement (Figure S1, S2). As a result of N170 being directed away from the active site, ceftazidime is able to bury more profoundly in the cavity, as exhibited by the docked complexes of acylated ceftazidime in the binding pocket of D179 variants. For orderly ceftazidime binding, omega loop restructuring that drives a disparate conformation of N170 appears necessary. This is also indicated by the MD simulations of ceftazidime acyl-enzyme complex with WT KPC-2.^[20]^ The resulting enhanced ceftazidime binding for KPC-2 variants is in agreement with kinetic studies that report reduced *K*_m_ values and propose that the elevated kinetics of ceftazidime binding is the focal cause for CAZ-AVI resistance.^[19,30,49]^ While the shift in N170 causes active site expansion and promotes increased binding affinity for the bulky cephalosporin, the deacylation mechanism, where N170 is involved in priming the water molecule, is compromised. Our findings are in accordance with kinetic and mass spectrometry assays that report slower ceftazidime hydrolysis/turnover by D179 variants compared to the WT β-lactamase. The catalytic turnover (*k*_cat_), as well as hydrolytic efficiency (*k*_cat_/*K*_m_), decreases for ceftazidime, owing to the reduced rate of deacylation. The “trapping” of β-lactams, as suggested by various kinetic studies, could be explained by the lowered rate of deacylation and slower hydrolysis by virtue of N170 displacement.^[19,27,29,37]^ However, the variants are still able to hydrolyze ceftazidime leading to overall increased MIC and drug resistance. This is unique to ceftazidime due to its bulky component, the aminothiazole ring on the C7 substituent, that is able to position in the pocket when N170 displaces away.^[29]^ In a broad sense, we propose that the altered susceptibility of KPC-2 variants to different β-lactam drugs is due to the active site expansion and N170 displacement. The loss of the stabilizing omega loop interactions results in a highly flexible structure unable to constrain N170 in its deacylation-competent orientation.

### D179 variants (D179N and D179Y) exhibit substrate-assisted catalysis for ceftazidime

We have demonstrated the shift in N170 as a probable explanation for the high binding affinity for ceftazidime but enhanced resistance to ceftazidime-avibactam is not a consequence of better drug entrapment solely. If the displaced conformation of N170 hinders deacylation, how is complete hydrolysis of ceftazidime achieved? Does N170 return to its native conformation after drug binding to deacylate the acyl-enzyme complex and regenerate the active enzyme? Or does some different mechanism come into play to mediate effective deacylation?

We speculate that with N170 oriented away from the active site, the ring amine or the primary amine of the aminothiazole group in ceftazidime is ideally positioned to activate the water molecule and initiate the attack on the covalent complex. To carry out the deacylation step of β-lactam hydrolysis, the positioning of a water molecule near the carbonyl of the acylated intermediate as well as a general base in the vicinity for hydrolytic water activation is crucial. In the absence of a base proximal to the active site in class-C β-lactamases, it was proposed that the β-lactam nitrogen, which is a secondary amine nitrogen post acylation and β-lactam ring opening, can activate the water molecule.^[50]^ We speculate that similar activation of the conserved water molecule is initiated via the aminothiazole group of ceftazidime in KPC-2 variants. Superimposition of the docked ceftazidime in the D179N active site with the WT revealed the distance between the water molecule and that of primary amine nitrogen and ring amine nitrogen to be 3.2 and 3.6 A°, respectively (Fig S3). Similarly, for D179Y, the respective distances are 3.8 and 4.1 A° (Fig S4). We posit that there is an “electrostatic sink” of the aminothiazole group where there is a possibility of an attack on the water molecule by two nitrogens within its vicinity. This way, the substrate itself promotes the approach of the deacylating water molecule to the acyl-enzyme complex.

### D179 variants show increased susceptibility to moxalactam

Moxalactam (MOX) is usually grouped with expanded spectrum cephalosporins, but the sulfur atom of the cephem nucleus is substituted with an oxygen atom (Fig S5). This synthetic oxacephem antibiotic shows potent antibacterial activity against ESBL-producing bacterial strains.^[51]^ While the kinetic studies suggesting its activity against KPC-producing strains are limited, *E.coli* harboring KPC-6 (V240G) reported elevated resistance to MOX with increased MIC.^[52]^ We questioned how the hydrolyzing profile of MOX is impacted by omega loop mutations in KPC at position 179. The emerging bacterial strains harboring KPC-2 substitutions show differential hydrolyzing activity/susceptibility for the two drugs (Table 1). We indicate that the structural changes around the active site, as a result of omega loop mutations, render the enzyme incapable to carry out substrate assisted deacylation with MOX. With the deacylation mechanism hindered, complete hydrolysis can be governed by peculiar structural characteristics of the substrate itself. For ceftazidime, though the deacylation mechanism is compromised due to N170 displacement, the positioning of the aminothiazole ring as a key structural moiety with two potential basic nitrogens can drive the deacylation of the carbonyl complex and lead to complete albeit slower β-lactam hydrolysis. On the other hand, KPC-2 mutants bearing D179 substitutions are likely to become susceptible to drugs like MOX, where the antibiotic structure lacks the aminothiazole ring to carry out the deacylation process. The absence of an aminothiazole ring or a similar structural feature completely eliminates the possibility of substrate-assisted catalysis, as observed for MOX. Thus, KPC-2 mutants lose their ability to hydrolyze MOX (confer resistance) as well as other β-lactams, and the antibacterial activity of this cephalosporin is restored. Mass spectrometry of MOX with D179Y showed a prolonged acyl-enzyme complex (Fig 6). There is fragmentation of MOX and only the 405 Da adduct is present. The proposed reaction mechanism for this fragmentation is provided (Fig S6).

**Figure 6:**
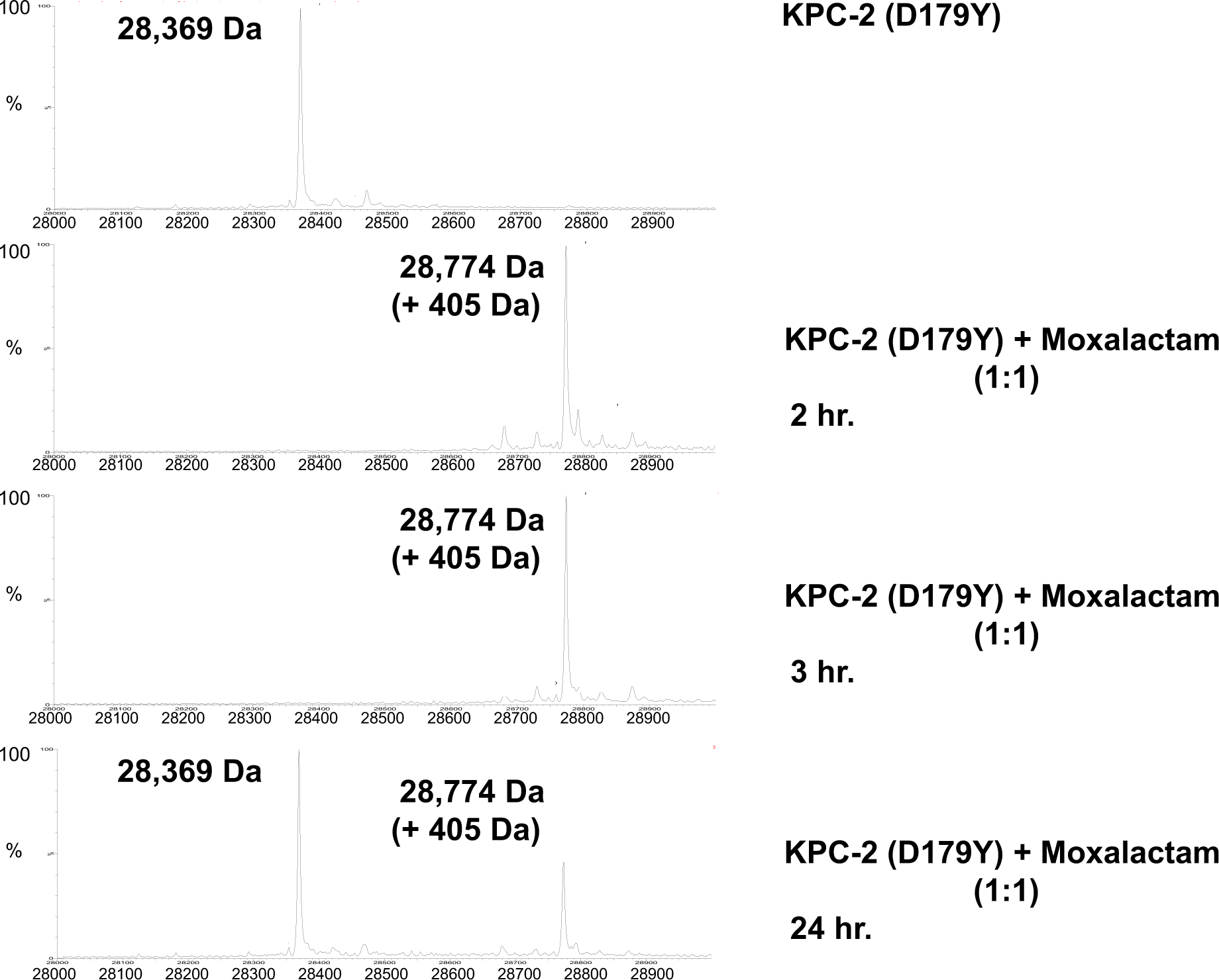
Timed Mass Spectrometry. The KPC-2 D179Y variant forms a prolonged (24 hr) acyl-complex with moxalactam.

**Table 1:**
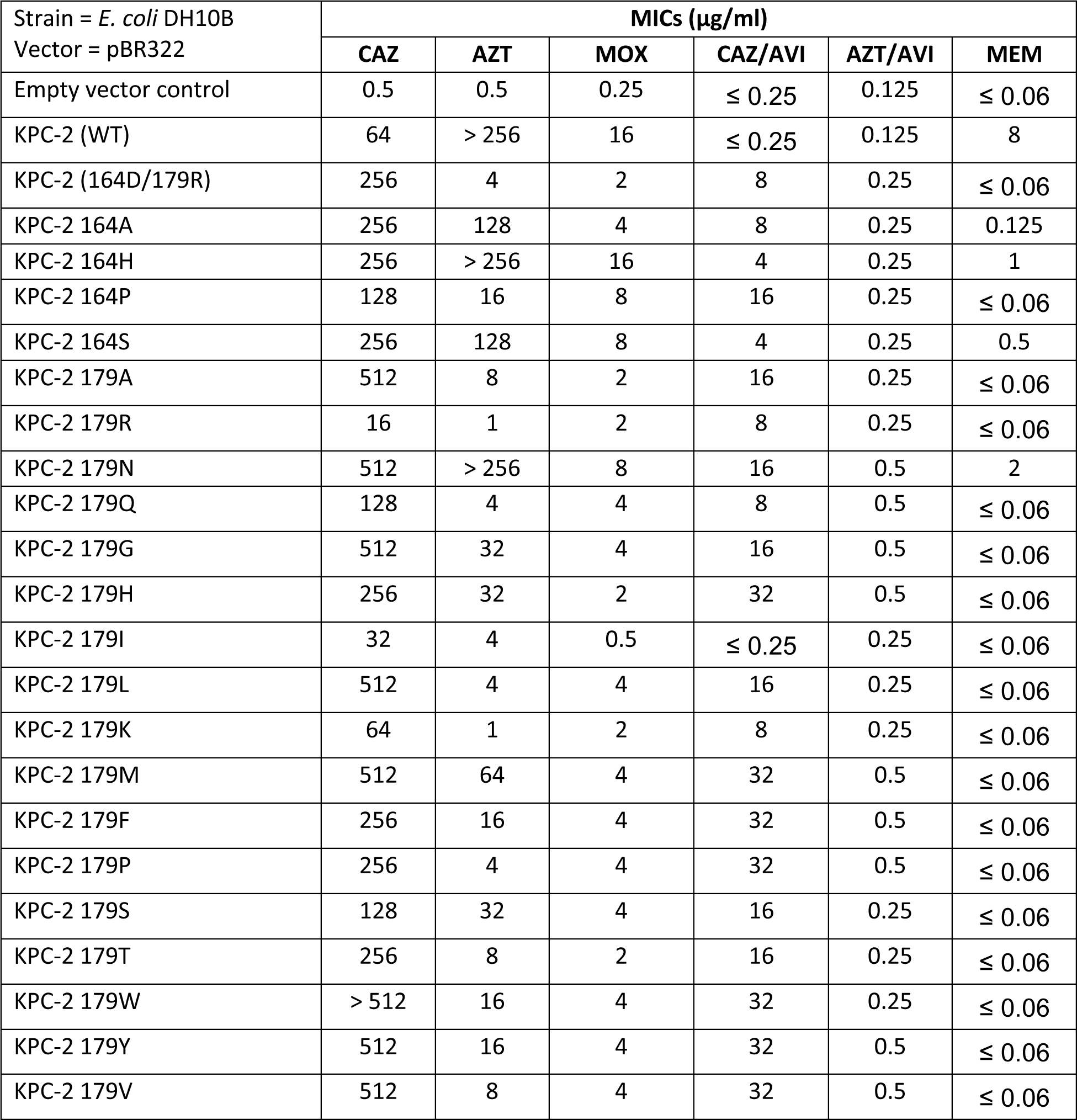
MICs of β-lactam and β-lactam–β-lactamase inhibitor combinations against KPC-2 D179 and R164 variants.

| Strain = *E. coli* DH10B<br>Vector = pBR322 | MICs ($\mu\text{g/ml}$) | | | | | |
| --- | --- | --- | --- | --- | --- | --- |
|  | CAZ | AZT | MOX | CAZ/AVI | AZT/AVI | MEM |
| Empty vector control | 0.5 | 0.5 | 0.25 | $\leq 0.25$ | 0.125 | $\leq 0.06$ |
| KPC-2 (WT) | 64 | > 256 | 16 | $\leq 0.25$ | 0.125 | 8 |
| KPC-2 (164D/179R) | 256 | 4 | 2 | 8 | 0.25 | $\leq 0.06$ |
| KPC-2 164A | 256 | 128 | 4 | 8 | 0.25 | 0.125 |
| KPC-2 164H | 256 | > 256 | 16 | 4 | 0.25 | 1 |
| KPC-2 164P | 128 | 16 | 8 | 16 | 0.25 | $\leq 0.06$ |
| KPC-2 164S | 256 | 128 | 8 | 4 | 0.25 | 0.5 |
| KPC-2 179A | 512 | 8 | 2 | 16 | 0.25 | $\leq 0.06$ |
| KPC-2 179R | 16 | 1 | 2 | 8 | 0.25 | $\leq 0.06$ |
| KPC-2 179N | 512 | > 256 | 8 | 16 | 0.5 | 2 |
| KPC-2 179Q | 128 | 4 | 4 | 8 | 0.5 | $\leq 0.06$ |
| KPC-2 179G | 512 | 32 | 4 | 16 | 0.5 | $\leq 0.06$ |
| KPC-2 179H | 256 | 32 | 2 | 32 | 0.5 | $\leq 0.06$ |
| KPC-2 179I | 32 | 4 | 0.5 | $\leq 0.25$ | 0.25 | $\leq 0.06$ |
| KPC-2 179L | 512 | 4 | 4 | 16 | 0.25 | $\leq 0.06$ |
| KPC-2 179K | 64 | 1 | 2 | 8 | 0.25 | $\leq 0.06$ |
| KPC-2 179M | 512 | 64 | 4 | 32 | 0.5 | $\leq 0.06$ |
| KPC-2 179F | 256 | 16 | 4 | 32 | 0.5 | $\leq 0.06$ |
| KPC-2 179P | 256 | 4 | 4 | 32 | 0.5 | $\leq 0.06$ |
| KPC-2 179S | 128 | 32 | 4 | 16 | 0.25 | $\leq 0.06$ |
| KPC-2 179T | 256 | 8 | 2 | 16 | 0.25 | $\leq 0.06$ |
| KPC-2 179W | > 512 | 16 | 4 | 32 | 0.25 | $\leq 0.06$ |
| KPC-2 179Y | 512 | 16 | 4 | 32 | 0.5 | $\leq 0.06$ |
| KPC-2 179V | 512 | 8 | 4 | 32 | 0.5 | $\leq 0.06$ |

### R164 mutants (R164S and R164H) also present a shift in N170 leading to CAZ-AVI resistance

To investigate the modified conformational landscape of R164S and R164H, we used the backbone dihedral angles of residue R164 to characterize the free energy surface. The free energy plots resulted in one basin for both R164S and R164H mutants. (Fig S7) The extracted conformations highlight the overall conserved omega loop structure, unlike D179 mutants, where the loop becomes disordered as a consequence of R164-D179 salt bridge disruption. Due to the substitution of arginine with serine and histidine in R164S and R164H KPC-2, respectively, the native interactions R164-D179, R164-S171, and R164-E168 are lost in the variant enzymes. However, as a result of the solvent-exposed conformation of R164, the substitution at this position doesn’t drastically perturb the orientation of other residues within or outside the loop. Our analysis shows that the fluctuation in the loop of R164 mutants is not as dynamic as in D179 mutants, but the E166-N170 interaction is still lost. Similar to D179 KPC variants, R164S and R164H KPC-2 also present a shift in the side chain of N170, where it moves out facing away from the active site. We propose that the altered enzymatic activity of KPC-2 bearing substitutions at 164 and 179 positions of the omega loop is due to the modified conformation of N170. With the side chain of N170 moved out, the mutant enzymes are primed to accommodate the bulky aminothiazole group of ceftazidime, resulting in enhanced kinetics of ceftazidime binding. This has also been suggested earlier by circular dichroism experiments for KPC-2 R164S mutant.^[24]^ The structural perturbation of N170 around the active site translates to the ceftazidime avibactam resistance phenotype. The lack of stabilizing salt bridge interaction at the neck of the omega loop drives N170 displacement and a consequent expanded binding pocket. However, the flexible omega loop renders the enzyme incapable of sustaining structural deformations required for the binding of several other drugs like ampicillin, cephalothin, cefotaxime, cefepime, aztreonam, and imipenem.^[24]^

The deacylation mechanism is also impaired, resulting in reduced MICs to these drugs. For KPC-2 omega loop mutants, we observe a different conformation of N170 in the apo-enzyme as a result of omega loop mutation, whereas for SHV-type β-lactamase, this structural disorder is only noticed upon binding of the ligand.^[40]^ Since there is a ligand-induced disorder, ceftazidime binding faces initial resistance due to potential steric clashes between the aminothiazole group and N170 facing the active site. Thus, *K*_m_ for ceftazidime slightly increases for SHV-1 β-lactamase, as opposed to KPC-2, where *K*_m_ decreases, and enhanced binding affinity for ceftazidime is identified as the resistant determinant.^[40]^

## Conclusions

In summary, the application of wt-MetaD simulations, microbiological, biochemical and crystallographic analysis of these omega loop variants enhance our understanding of this extremely problematic phenotype resulting in ceftazidime avibactam resistance. In terms of future drug development, the underappreciated role of the aminothiazole group in substrate assisted catalysis is a starting point for developing novel β-lactams. The unexpected increase in susceptibility to the representative oxacephem suggests a path forward in the design of novel cephems. Further studies exploring the interactions of carbapenems or carbapenem mimics in these variants will also further illuminate their potential role as carbapenemase inhibitors.

## Methods

### Mass spectrometry

Ten micrograms of *bla*_KPC-2(D179Y)_ were incubated with the substrate (MOX) at a molar ratio of 1:1 in sterile 10 mM phosphate-buffered saline (PBS) at pH 7.4 for a total reaction volume of 20 µl for the times indicated in the figures. Reactions were quenched with 1% acetonitrile and 0.1% formic acid. Samples were analyzed using Q-TOF Waters Synapt-G2-Si and a Waters Acquity ultrapressure liquid chromatography (UPLC) BEH C18 column (1.7-µm pore size; 2.1 by 50 mm). MassLynxV4.1 was used to deconvolute protein peaks. The tune settings for each data run were as follows: capillary voltage at 3.5 kV, sampling cone at 35, source offset at 35, source temperature of 100°C, desolvation temperature of 500°C, cone gas at 100 liters/h, desolvation gas at 800 liters/h, and nebulizer bar at 6.0. Mobile phase A was 0.1% formic acid–water. Mobile phase B was 0.1% formic acid–acetonitrile. The mass accuracy for this system is ±5 Da.

### Minimum inhibitory concentration (MIC) measurements

MIC measurements were performed in triplicate using the Mueller-Hinton (M-H) agar dilution method, according to the Clinical Laboratory and Standards Institute (CLSI) protocol.^[53]^ Briefly, bacterial cultures containing *bla*_KPC_ variants previously cloned into a uniform vector (pBR322)^24,27^ were incubated overnight at 37 °C in M-H broth. The cultures were diluted, and an Oxoid replicator was used to deliver 10 μL of a culture containing approximately 10^4^ CFU. MICs were determined for the following: ceftazidime (CAZ), aztreonam (AZT), moxalactam (MOX), ceftazidime/avibactam (CAZ/AVI), aztreonam/avibactam (AZT/AVI), and meropenem (MEM).

### Computational methods

The structure of KPC-2 WT (PDB id-3DW0) was obtained from the RCSB protein data bank. The ICM mutagenesis program was employed for the *in silico* construction of the six variants (D179N, D179Y, D179Q, D179A, R164S, and R164H).^[54]^ The system was prepared using a high-throughput molecular dynamics (HTMD) protocol.^[55]^ One covalent disulphide bond was added between Cys 69 and 238. The protein parameters were defined using the Amberff14SB force field,^[56,57]^ and explicit TIP3P water molecules were included in a cubic box with its edge at least 10 Å from the closest solute atom.^[58]^ The systems were generated using 0.15M NaCl. The electrostatic interaction distances were set to ≤ 8 Å. Long range electrostatic interactions were calculated using the particle mesh Ewald summation. The systems underwent a minimization process with 1,000 steps of steepest descent integrator, followed by a 5 ns equilibration in the NPT ensemble with a Berendsen barostat at 1 atm.^[59]^ The temperature was maintained at 300 K using a Langevin thermostat. For all systems, a 3 μs production run was conducted in the NVT ensemble with a time step of 4 fs, utilizing the ACEMD program.^[60]^ To enhance sampling, Well-Tempered Metadynamics simulations (wt-MetaD) were performed at 300 K using the equilibrated structures of both the wild-type KPC-2 and variant structures.^[39,42]^ These simulations were carried out using the ACEMD software and the PLUMED 1.3 plug-in, with an integration step of 4 fs.

To gain insight into how key mutations impact the omega loop dynamics, we examined the changes in the free energy landscape with respect to backbone dihedral angles of the WT/variant residue. These dihedral angles are particularly relevant as they capture the underlying slowest motions and reflect the structural effects of substitutions, thus governing our choice of collective variables (CV1 = ϕ and CV2 = ϕ of the substituted residue).^[44]^ To introduce a bias, Gaussian distributions with widths of 0.1 and 0.1 radians were applied to the two collective variables (CVs), respectively, while the height of the Gaussians was set at 0.5 kJ/mol. The Gaussians were deposited every 4 ps, ensuring a deposition rate of 0.125 kJ/(mol·ps). The bias factor remained constant at 15. Initially, we ran our simulations for 1.5 µs and then increased to 3 µs to achieve maximum convergence of the free energy landscape (Fig S8-S9). We observed our system to be diffusing rapidly in the entire CV space and visiting the relevant conformations with 3 µs of sampling time in the NVT ensemble. From the wt-MetaD trajectories, structures associated with the minima were identified based on the values of the collective variables CV1 and CV2 and were used to analyze the altered H-bonding network. To extract representative structures that capture the essence of each minimum, the conformations from the 300 K replica were clustered using Cα RMSD.

In addition to the wt-MetaD simulations, classical equilibrium MD simulations were run on the extracted representative conformations from each of the basins. The unbiased simulations were run in duplicates for 1 μs each (Table S1). The analyses focused on the conformational drift (RMSD) of the enzyme (Fig S10-S11) and the sampling of the χ1 angle of N170 (Fig S12-17). The simulation protocol and analysis has been detailed in the supporting information.

Antibiotics ceftazidime and moxalactam were covalently docked to the representative structures from the free energy minima using the ICM-Pro software.^[54]^ Flexible ligand docking with the ICM-Pro software uses Monte Carlo simulations to globally optimise a set of internal ligand coordinates in the space of grid potential maps calculated for the protein catalytic site. The automatically determined Monte Carlo run length was extended by a multiplier (thoroughness) of ten, and five docking conformations were obtained. The structural figures were generated using Protein Imager (https://3dproteinimaging.com), a modified version of open-source PyMol (github.com/bieniekmateusz/pymol-mdanalysis), and Molsoft ICM-Pro package (http://www.molsoft.com).

## Supporting Information

Supporting information contains Figures S1-S17, Table S1

## Author Contribution

Simulations – Diksha Parwana, Jing Gu, Qinxin Wang; Mass Spectrophotometry – Chris Bethel, Emma Marshall, Andrea Hujer, MIC experiments - Chris R Bethel, Emma Marshall, Andrea M Hujer, Supervision – Robert A Bonomo and Shozeb Haider. All authors contributed in the preparation of the manuscript.

## Data Availability

All simulation data can be downloaded from https://doi.org/10.5281/zenodo.8433386

## Acknowledgements

This work was supported in part by funds and/or facilities provided by the Cleveland Department of Veterans Affairs, Award Number 1I01BX001974 (R.A.B.), from the Biomedical Laboratory Research & Development Service of the VA Office of Research and Development, and the Geriatric Research Education and Clinical Center VISN 10. The content is solely the authors’ responsibility and does not necessarily represent the official views of the NIH or the Department of Veterans Affairs. RAB is also supported by Merch, VenatoRx, Entasis, Shionogi, Wockhardt for research support only. All other authors declare no conflict of interest.

