## Supplementary for "The Structural Role of N170 in Substrate-assisted Deacylation in KPC-2 β-Lactamase"

**SUPPORTING INFORMATION**

[1]UCL School of Pharmacy;

[2]Research Service, Louis Stokes Cleveland Department of Veterans Affairs Medical Center, Cleveland, OH, USA; [3]Department of Medicine, Case Western Reserve University School of Medicine, Cleveland, OH, USA; [4]Clinician Scientist Investigator, Louis Stokes Cleveland Department of Veterans Affairs Medical Center, Cleveland, OH, USA; Department of Molecular Biology and Microbiology, Pharmacology, Biochemistry, and Proteomics and Bioinformatics, Case Western Reserve University School of Medicine, Cleveland, OH, USA; and CWRU-Cleveland VAMC Center for Antimicrobial Resistance and Epidemiology (Case VA CARES) Cleveland, OH, USA; [5]UCL Centre for Advanced Research Computing, London, UK.

Correspondence:

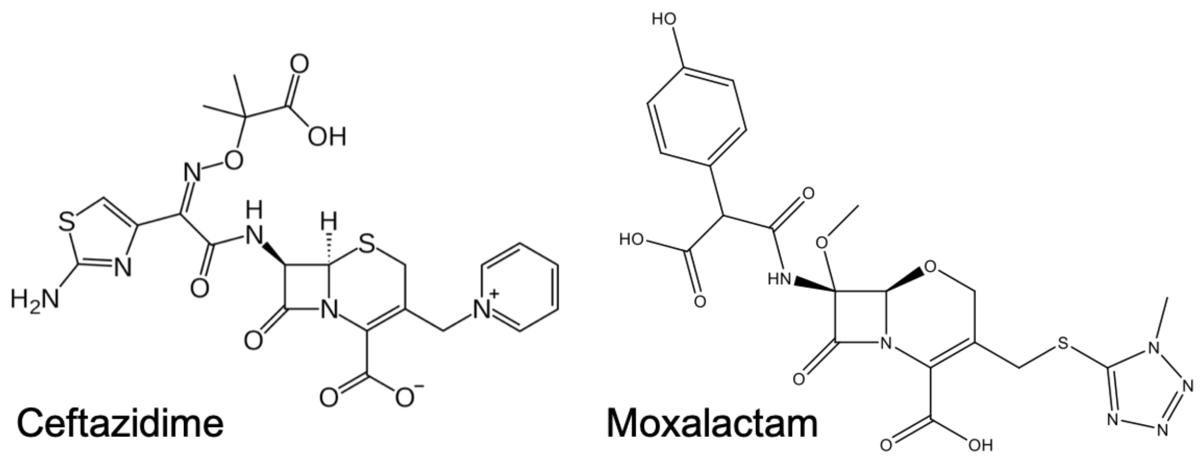


**Figure S1:** Structures of ceftazidime and moxalactam.

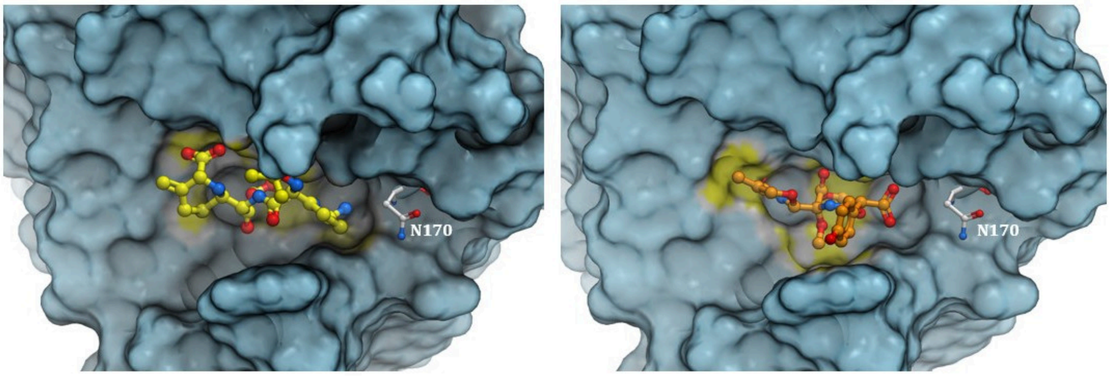


**Figure S2:** Covalent docking of Ceftazidime (left) and Moxalactam (right) in basin A of KPC-2 D179N

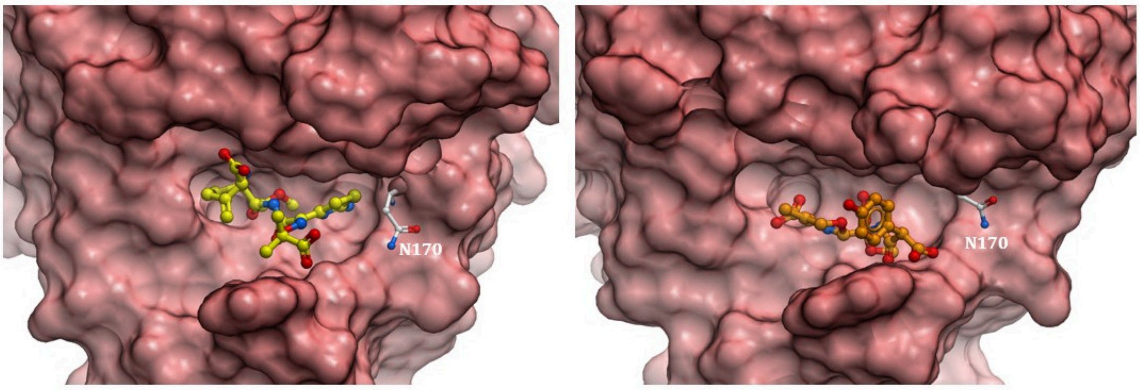


**Figure S3:** Covalent docking of Ceftazidime (left) and Moxalactam (right) in basin A and basin B of KPC-2 D179Y

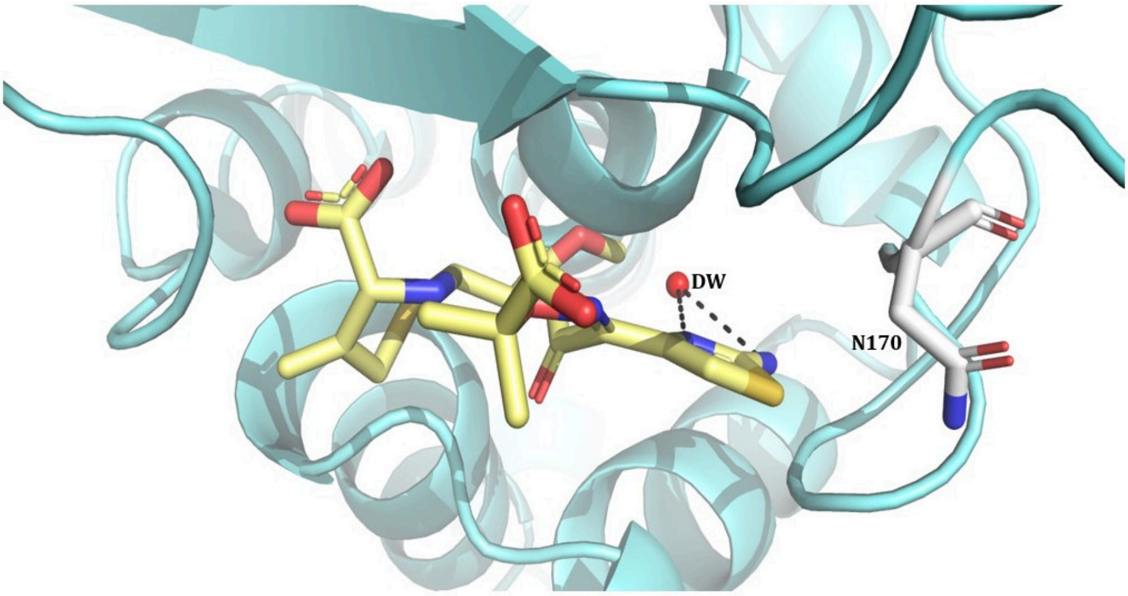


**Figure S4:** Covalent docking of ceftazidime to KPC-2 D179N. Interactions of the deacylating water molecule (DW) with triazole-ring nitrogen (3.7 Å) and primary amine (4 Å) of the aminothiazole group of ceftazidime are illustrated.

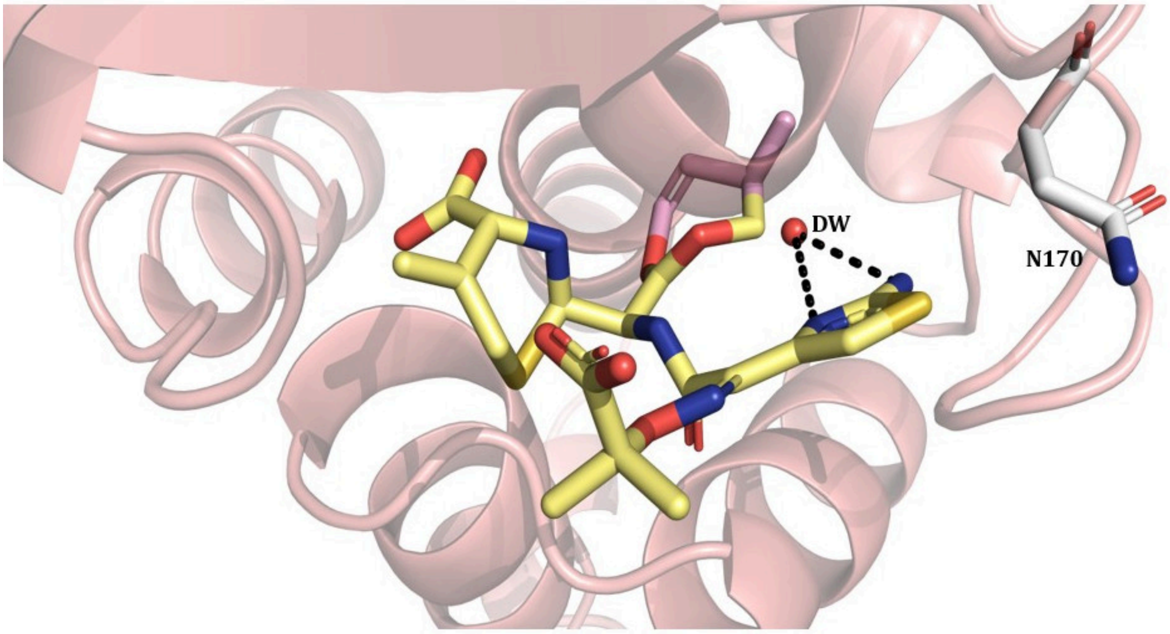


**Figure S5:** Covalent docking of ceftazidime to KPC-2 D179N. Interactions of the deacylating water molecule (DW) with triazole-ring nitrogen (3.9 Å) and primary amine (4.2 Å) of the aminothiazole group of ceftazidime are illustrated.

**Figure S6-** Proposed reaction mechanism for *bla*$_{KPC\text{-}2(D179Y)}$:moxalactam acyl-enzyme formation. Loss of the **R** group results in a prolonged covalent species of mass +405 Da as shown by mass spectrometry.

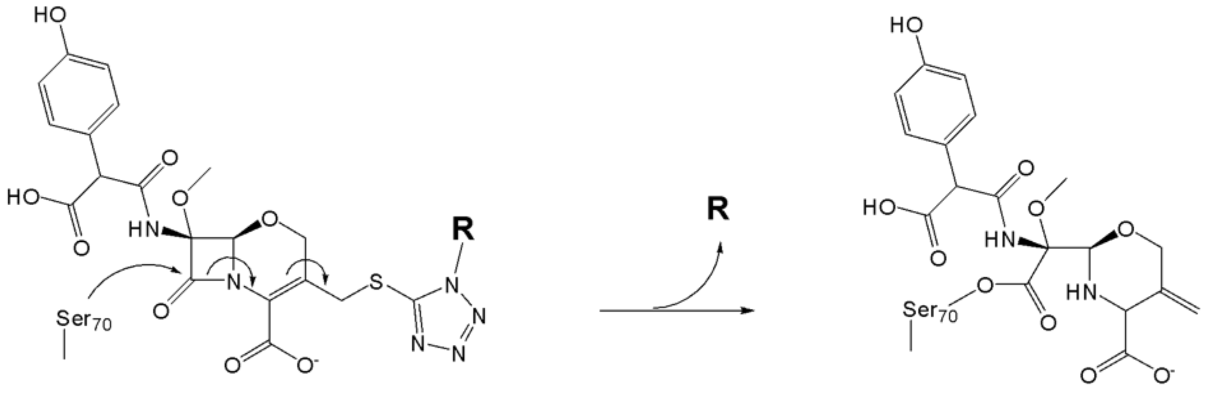

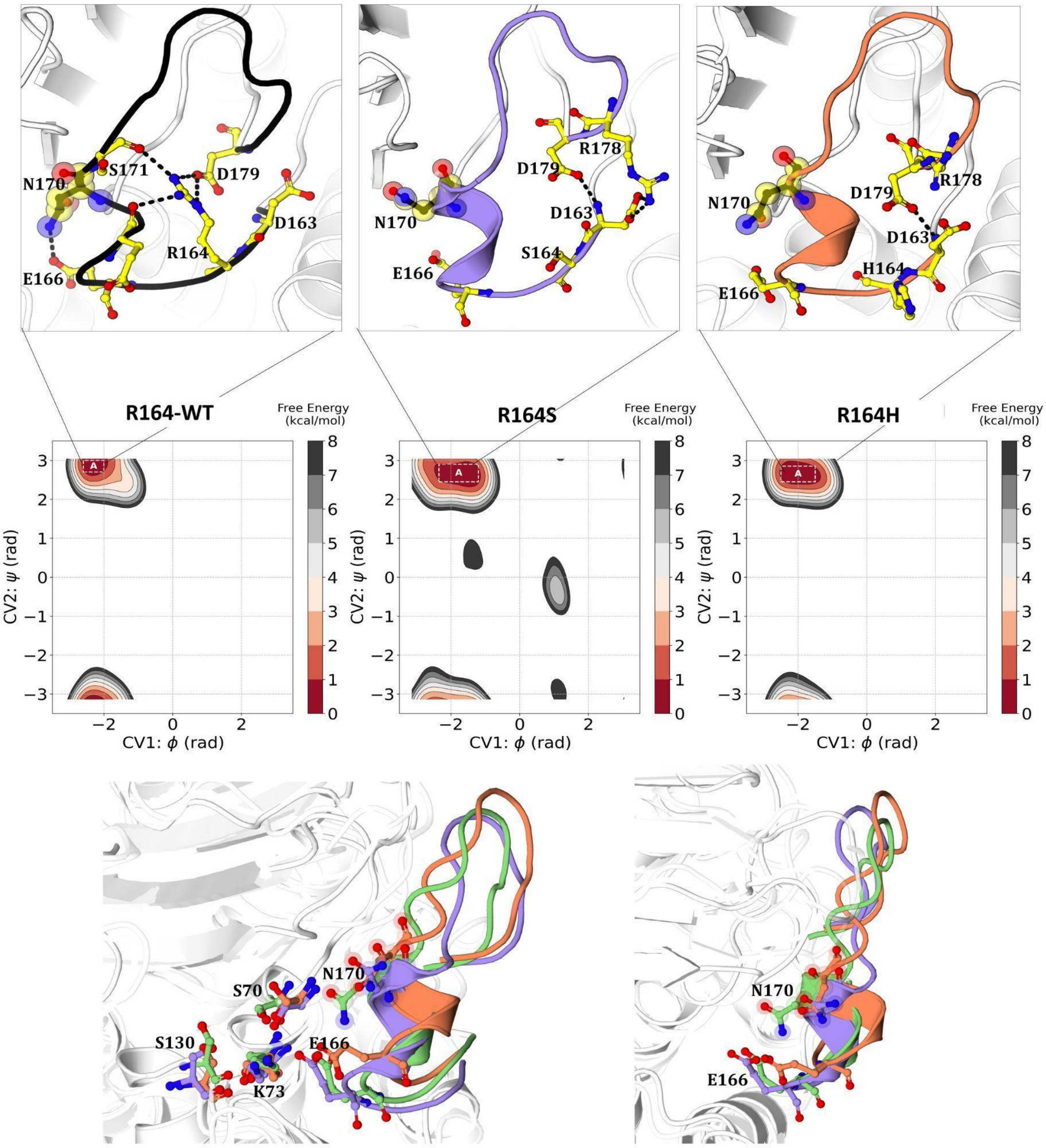


**Figure S7**- (A) Free energy plot for KPC-2 WT, R164S, and R164H as a function of φ (CV1) and ψ (CV2) dihedral angles of residue 164. Omega loop interactions of the conformations extracted from their corresponding free energy minima have been illustrated. (B) Superimposition of the extracted structures from free energy basins with the crystal structure of KPC-2 WT (PDB id-3dw0).

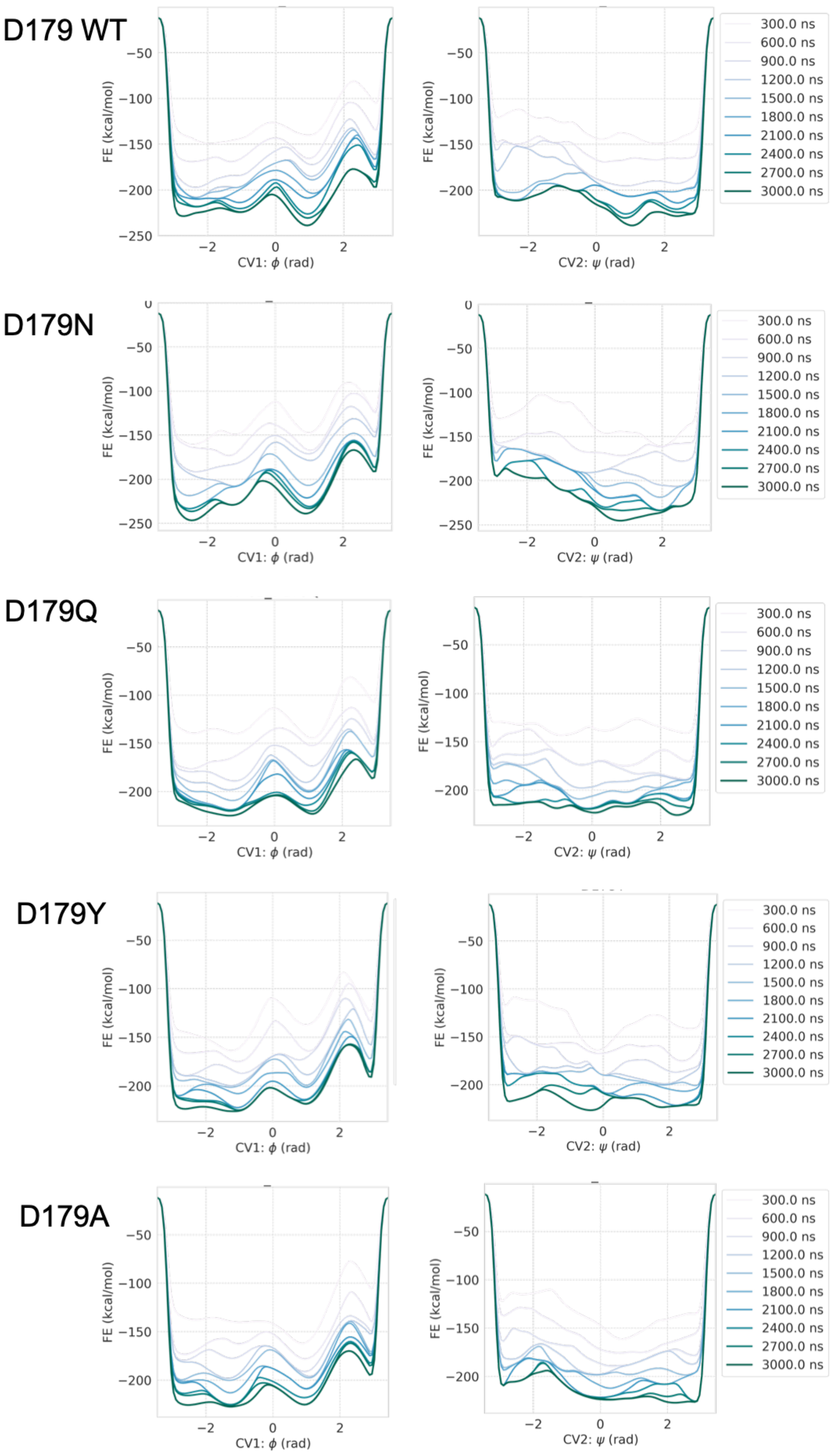


**Figure S8**- Convergence plots for CV1 ϕ and CV2 ψ from D179 wild type and variant well-tempered MetaD simulations

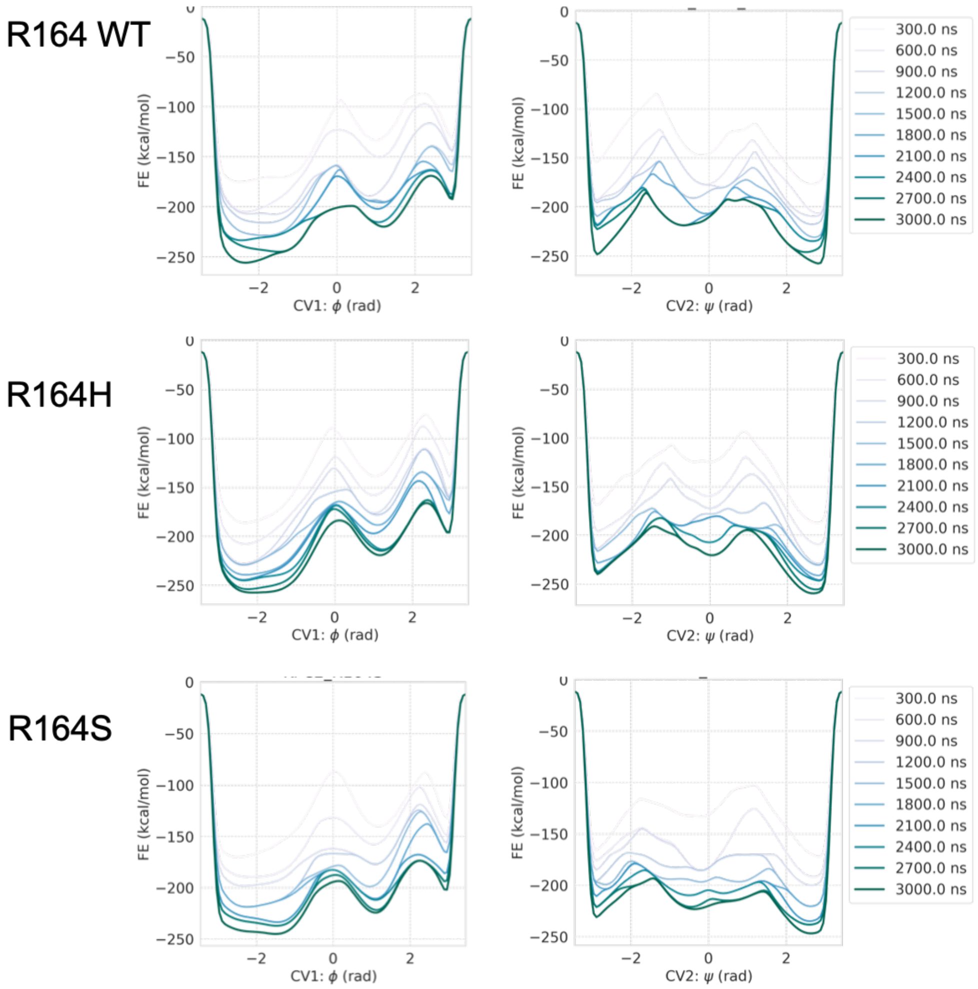


**Figure S9**- Convergence plots for CV1 $\phi$ and CV2 $\psi$ from R164 wild type and variant well-tempered MetaD simulations

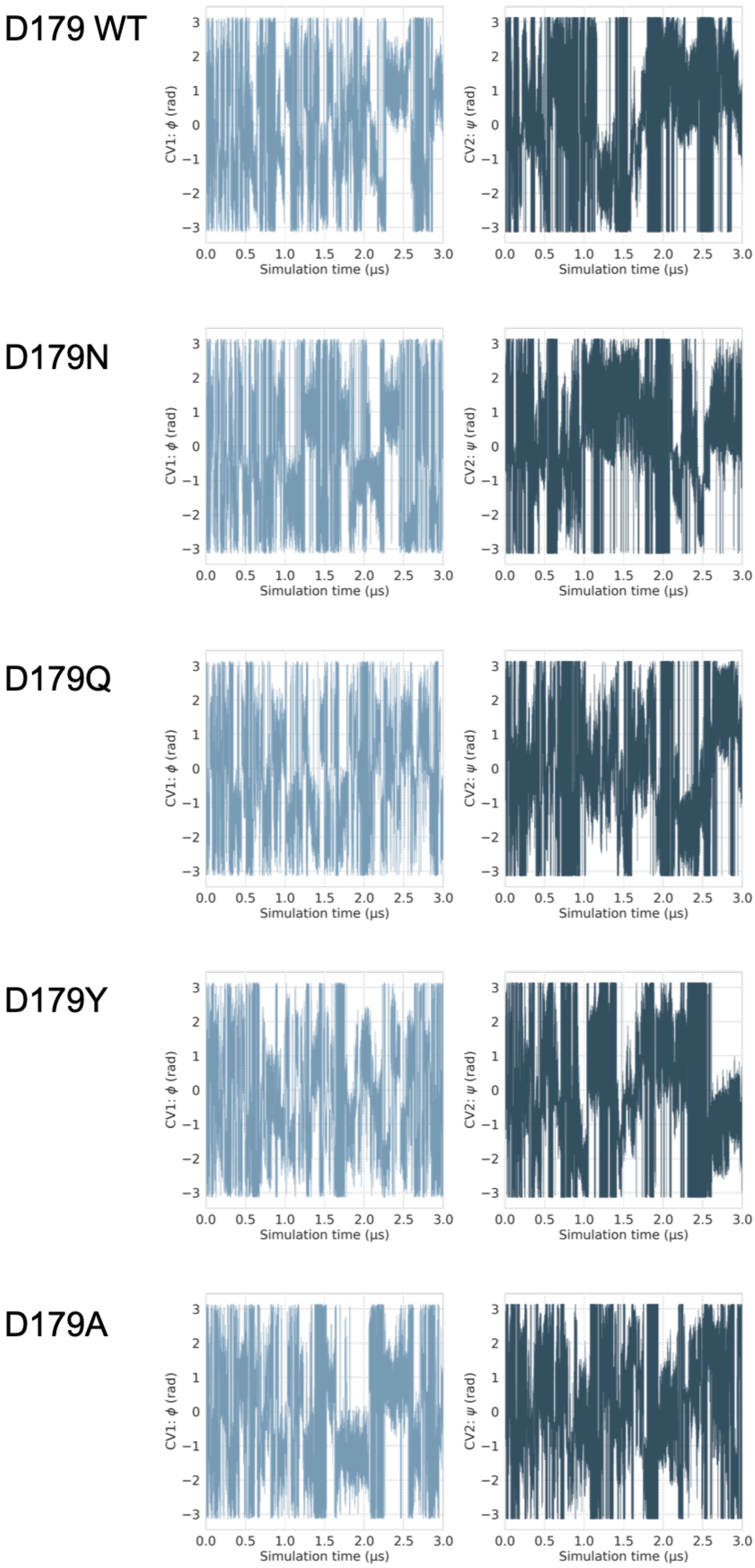


**Figure S10**- Diffusive behavior of D179 systems along the CVs ϕ and ψ in well-tempered MetaD simulations

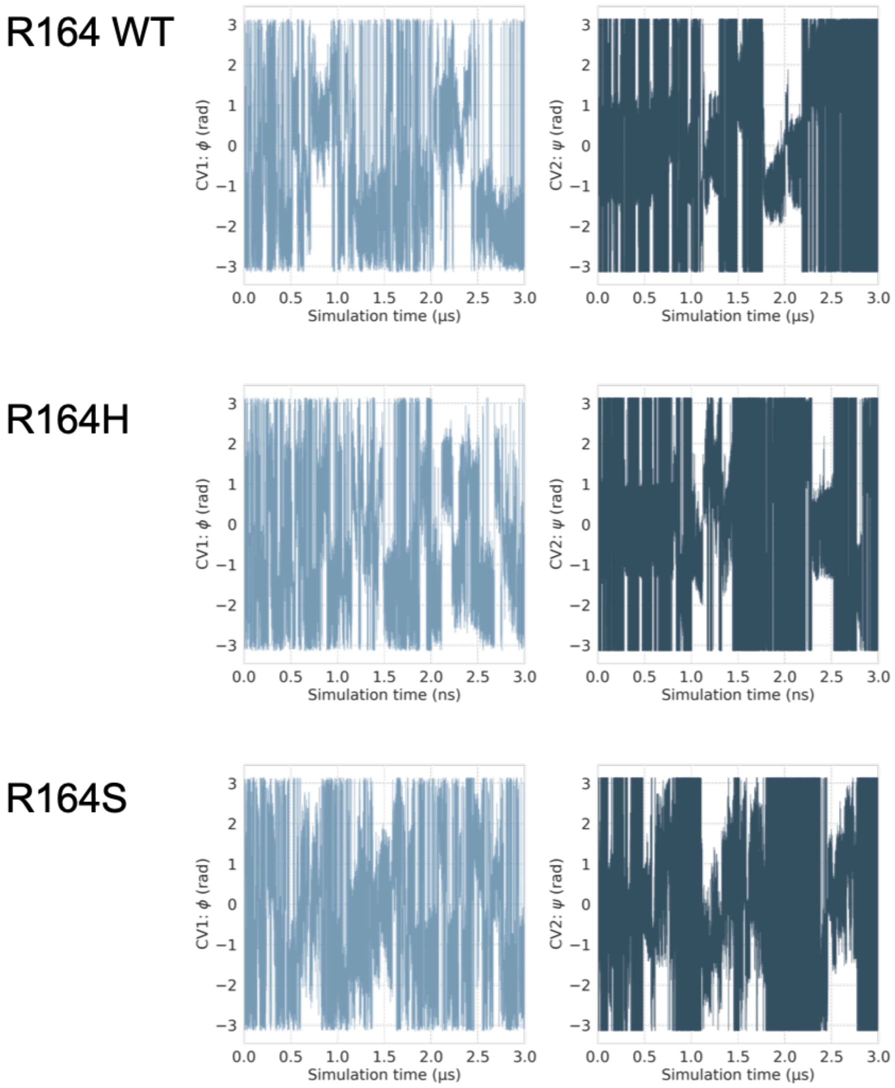


**Figure S11**- Diffusive behavior of R164 systems along the CVs $\phi$ and $\psi$ in well-tempered MetaD simulations

**Classical equilibrium molecular dynamics**

In addition to the wt-MetaD simulations, classical equilibrium simulations were run on the extracted representative conformations from each of the basins. The simulation protocol was exactly the same as for wt-MetaD simulations except for adding bias using plumed.[1] For the simulations, the structures were protonated at pH 7.0 using propKa as implemented in the playmolecule.[2] One covalent disulphide bond was added between Cys 69 and 238. The structures were then solvated in TIP3P water present in a cubic water box, whose edges were at least 10 Å from the closest solute atom.[3] Amber ff14SB force field was used to describe the protein atoms.[4] The electrostatic interaction distances were set to ≤ 8 Å. The long-range electrostatic interactions were computed using the particle mesh Ewald summation method.[3] Each system was then energy minimized using a steepest descent for 1000 steps. This was followed by an equilibration step for 5 ns at 1 atm using Berendsen barostat.[5] The initial velocities within each simulation were sampled from the Boltzmann distribution at 300K. Two replicates of simulations were run for each system for 1 μs each using the ACEMD molecular dynamics engine.[6] The production run was carried out under NVT ensemble using a Langevin thermostat with a damping of 0.1 $ps^{-1}$ and hydrogen mass repartitioning scheme to achieve a time step of 4 fs. Each trajectory frame was saved at 100 ps.

Table S1: Summary of classical molecular dynamics simulations.

| System | Basin | No. of Atoms | Simulation time | Average RMSD (nm) |
|---|---|---|---|---|
| WT – D179 | A | 27245 | 2 x 1 μs | 0.118 ± 0.009 |
| WT – R164 | A | 27245 | 2 x 1 μs | 0.117 ± 0.010 |
| D179Y | A | 26916 | 2 x 1 μs | 0.108 ± 0.008 |
| | B | 26916 | 2 x 1 μs | 0.133 ± 0.014 |
| | C | 26916 | 2 x 1 μs | 0.124 ± 0.013 |
| D179A | A | 26908 | 2 x 1 μs | 0.142 ± 0.012 |
| | B | 26908 | 2 x 1 μs | 0.135 ± 0.013 |
| | C | 26908 | 2 x 1 μs | 0.146 ± 0.012 |
| | D | 26908 | 2 x 1 μs | 0.132 ± 0.017 |
| D179Q | A | 26912 | 2 x 1 μs | 0.141 ± 0.009 |
| | B | 26912 | 2 x 1 μs | 0.116 ± 0.011 |
| | C | 26912 | 2 x 1 μs | 0.114 ± 0.013 |
| D179N | A | 26909 | 2 x 1 μs | 0.107 ± 0.011 |
| | B | 26909 | 2 x 1 μs | 0.125 ± 0.016 |
| R164H | A | 27237 | 2 x 1 μs | 0.133 ± 0.018 |
| R164S | A | 27231 | 2 x 1 μs | 0.150 ± 0.028 |
| | **Total** | | **32 μs** | |

The RMSD analysis (Table S1, Fig S12-S13) of all the systems simulated in the classical equilibrium molecular dynamics indicated that there was no untoward instability observed throughout the simulations. The main focus for the equilibrium molecular dynamics was the conformational sampling of residue N170. The $\chi 1$ angle of N170 was monitored throughout the simulations. An ensemble of structural conformations of N170 were identified to highlight the orientation of the side chain (Fig S14-19). The N170 side chain is able to sample multiple conformations in all D179 variant simulations irrespective of which system it has been extracted from (Fig S14-18). This effect is less pronounced in the R164 variant simulations, consistent with experimental data (Fig S19). It is worth emphasizing that while wt-MetaD simulations focused on sampling the backbone dihedral angles of residues R164 and D179, the side chain of N170 displayed extensive conformational flexibility independently. This flexibility is retained and is also observed in equilibrium molecular dynamics simulations.

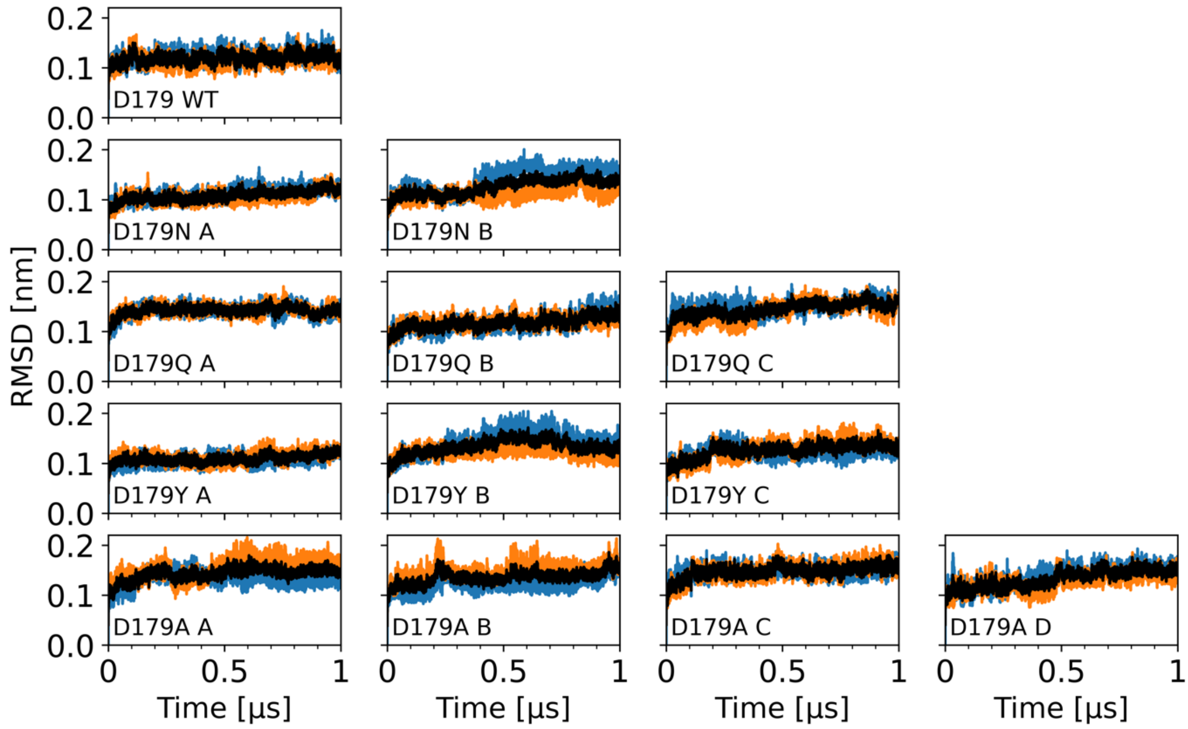


**Figure S12**: RMSD plots from D179 WT and variant classical unbiased MD simulations. The suffix alphabet denotes the basin from which the representative structure was extracted. The orange/blue lines are RMSD from duplicate simulations. The black line is the average of the two simulations.

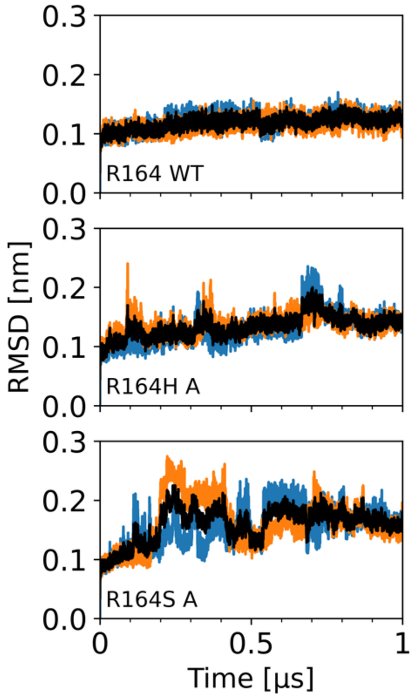


**Figure S13**: RMSD plots from R164 WT and variant classical unbiased MD simulations. The suffix alphabet denotes the basin from which the representative structure was extracted. The orange/blue lines are RMSD from duplicate simulations. The black line is the average of the two simulations.

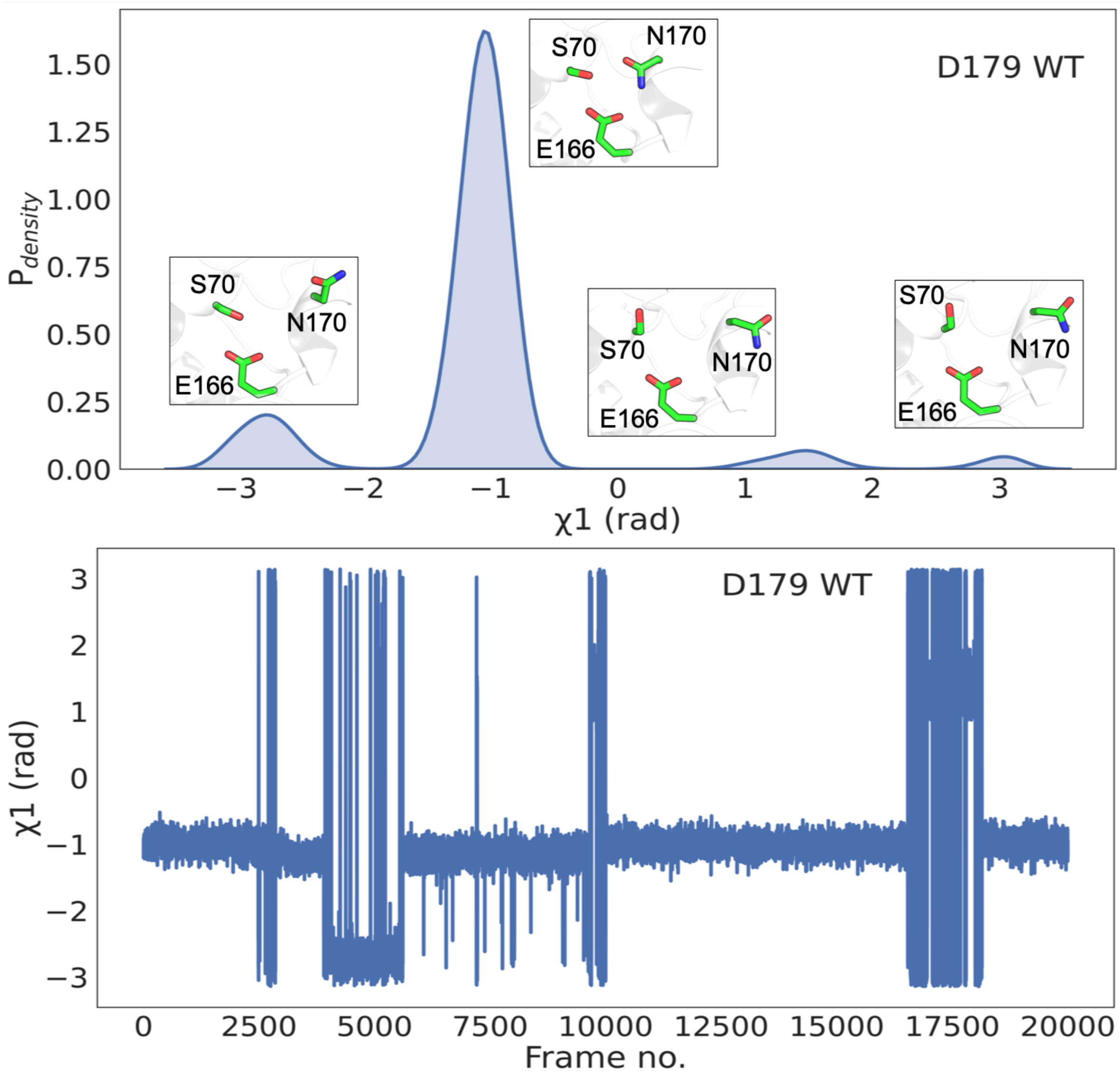


**Figure S14**: The conformations of the $\chi$1 dihedral angle of N170 in the D179 WT, calculated from the equilibrium simulations. The inset figures (top) illustrates the corresponding orientation of the N170, S70 and E166 side chains relative to N170.

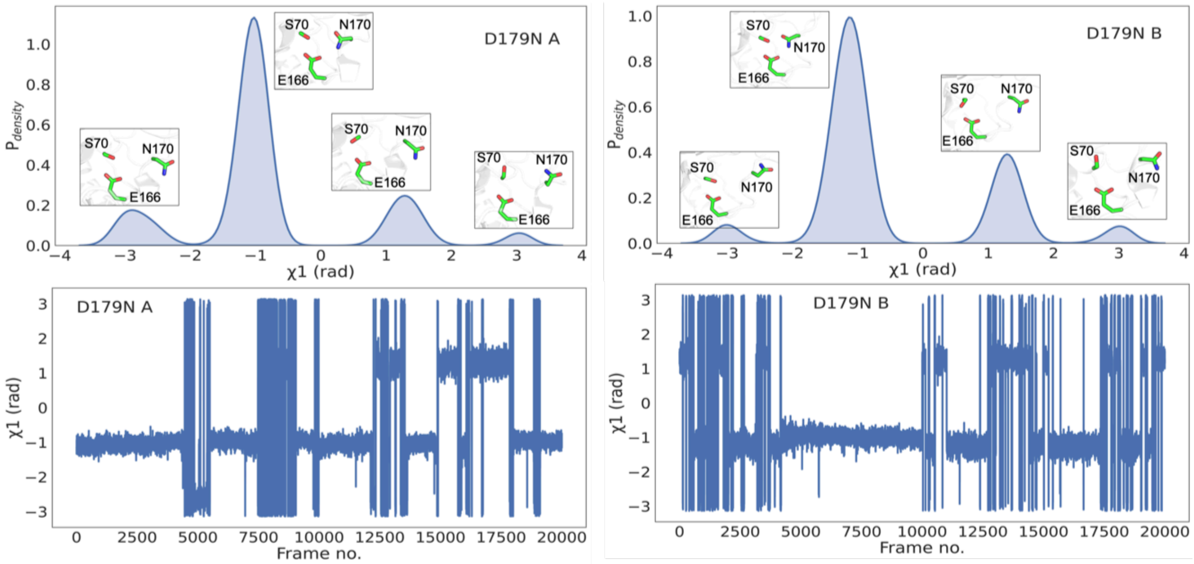


**Figure S15**: The conformations of the $\chi 1$ dihedral angle of N170 in the D179N variant, calculated from the equilibrium simulations. The inset figures (top) illustrates the corresponding orientation of the N170, S70 and E166 side chains relative to N170. System A and B are representative D179Y conformations extracted from the corresponding basins in the wt-MetaD simulations.

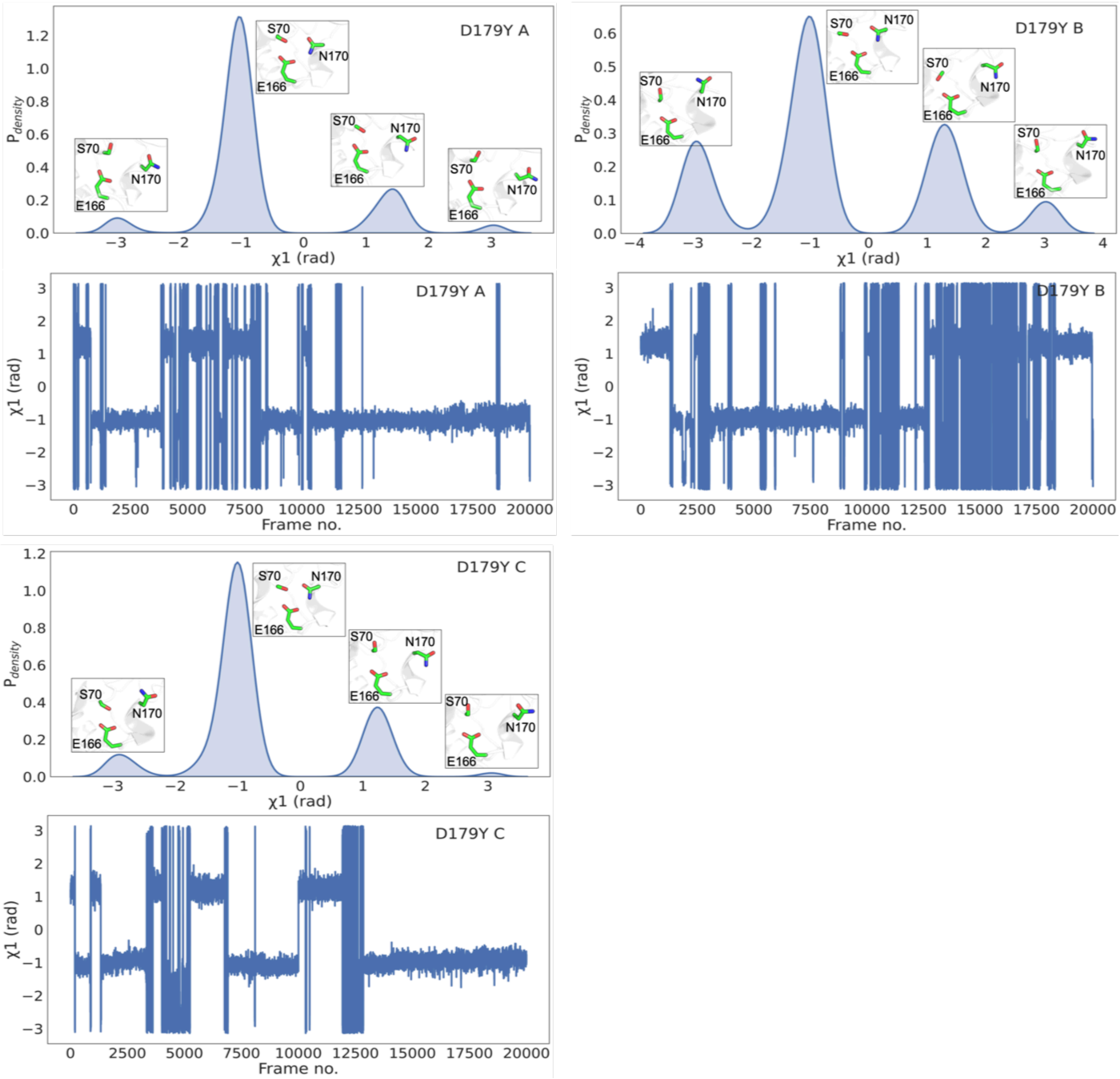


**Figure S16**: The conformations of the $\chi 1$ dihedral angle of N170 in the D179Y variant, calculated from the equilibrium simulations. The inset figures (top) illustrates the corresponding orientation of the N170, S70 and E166 side chains relative to N170. System A B, and C are representative D179Y conformations extracted from the corresponding basins in the wt-MetaD simulations.

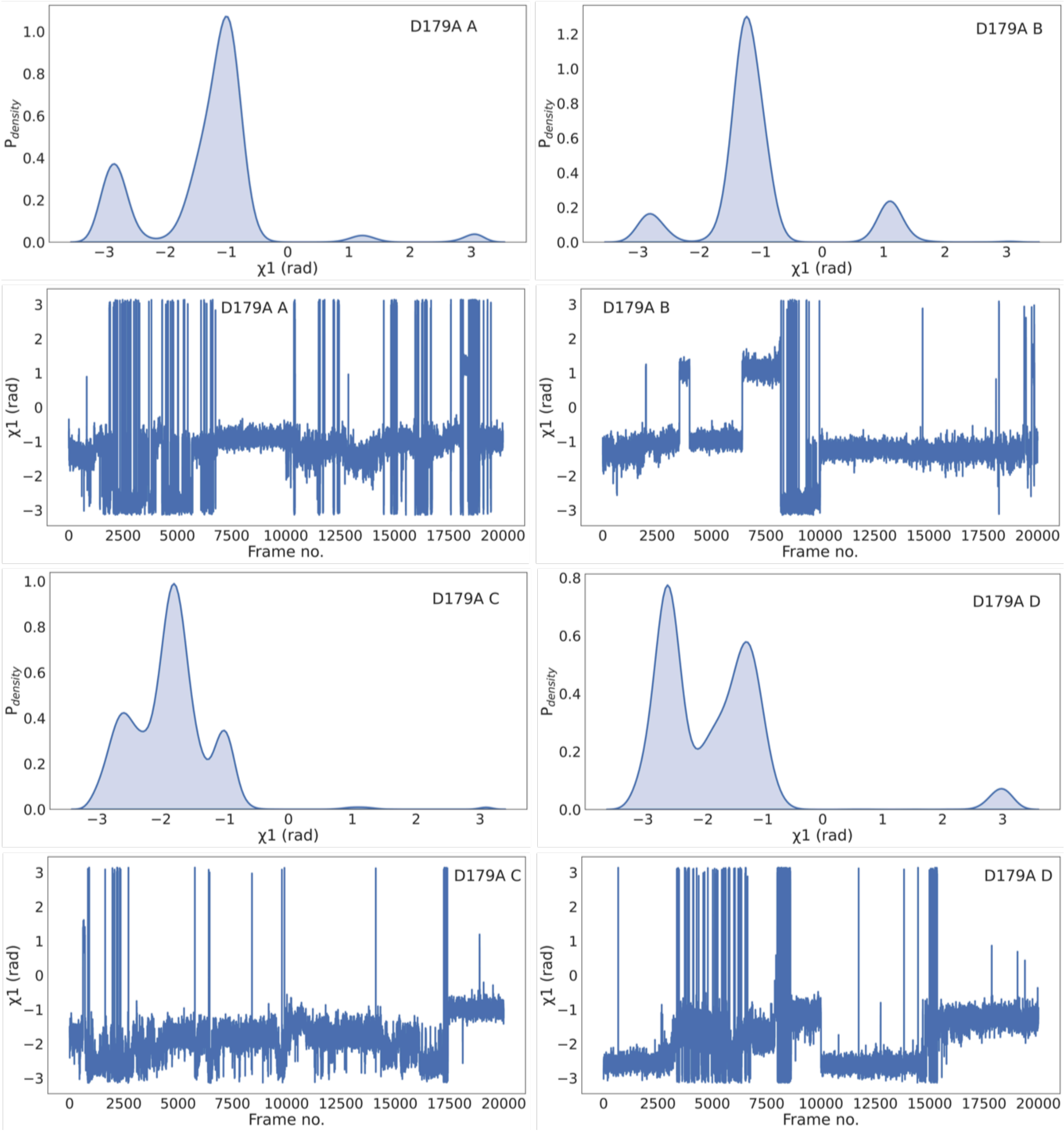


**Figure S17**: The conformations of the $\chi1$ dihedral angle of N170 in the D179A variant, calculated from the equilibrium simulations. System A B, C, and D is representative of D179A conformations extracted from the corresponding basins in the wt-MetaD simulations.

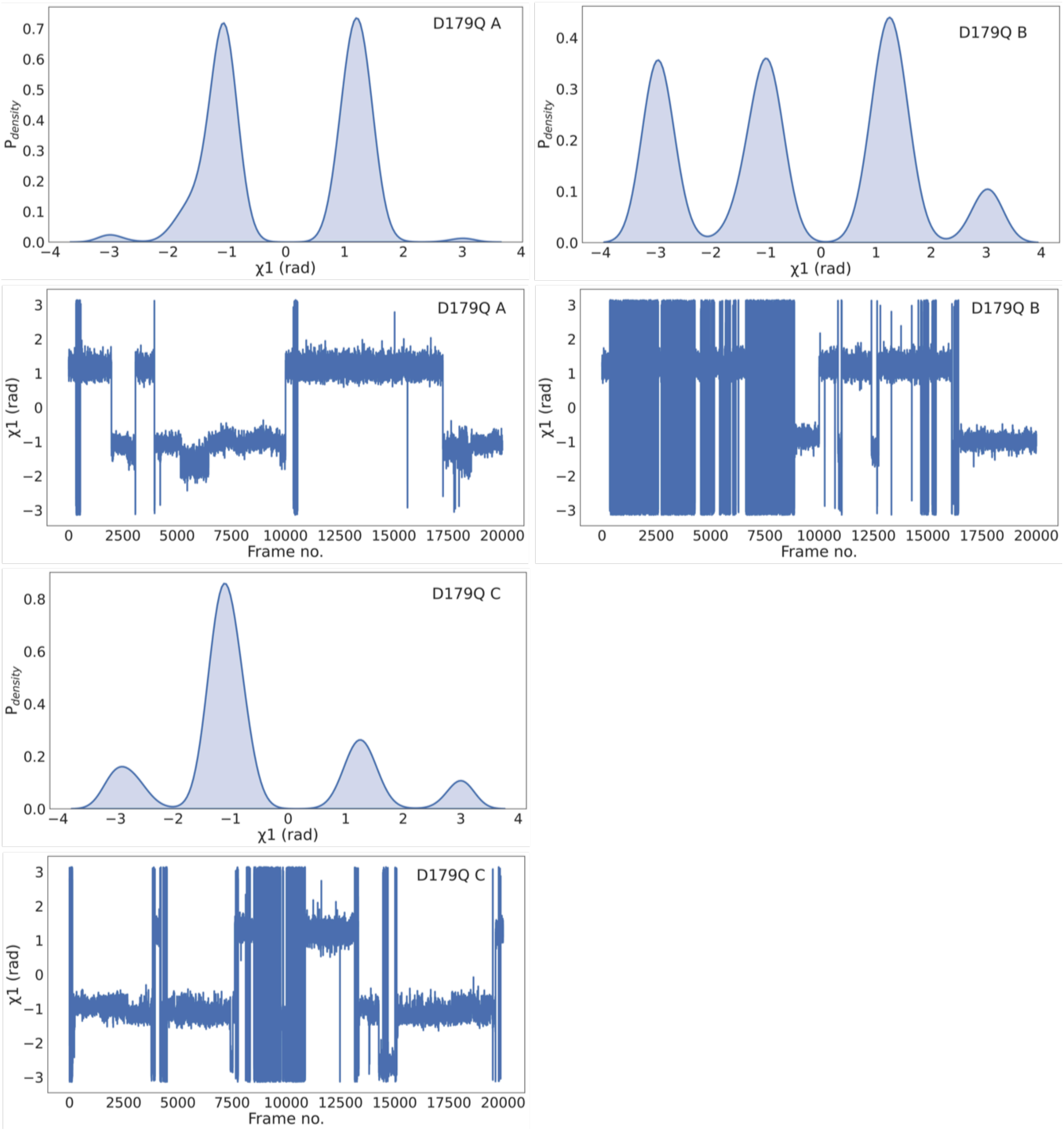


**Figure S18**: The conformations of the $\chi$1 dihedral angle of N170 in the D179Q variant, calculated from the equilibrium simulations. System A B, C, and D is representative of D179Q conformations extracted from the corresponding basins in the wt-MetaD simulations.

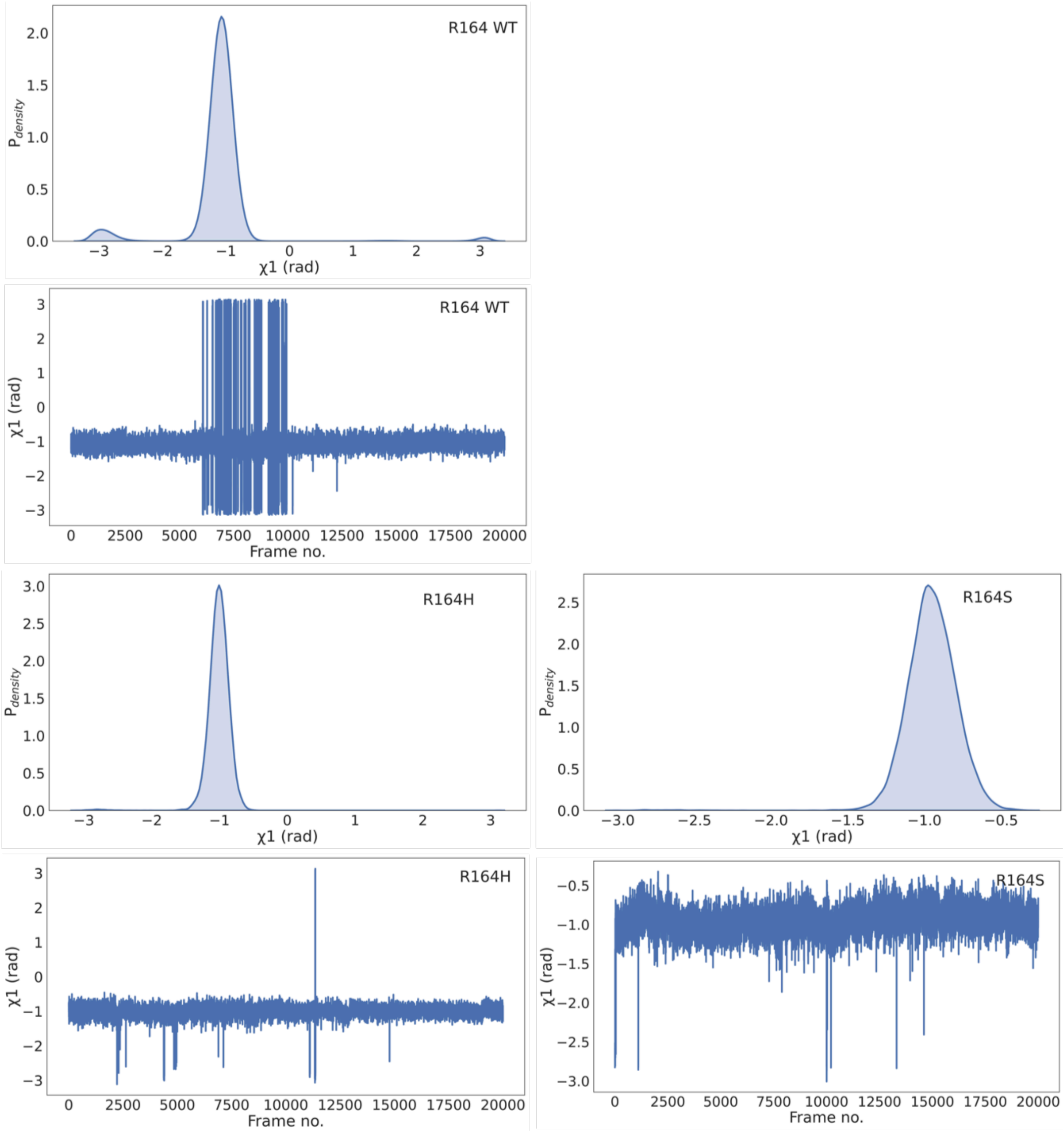


**Figure S19**- The conformations of the $\chi$1 dihedral angle of N170 in the R164 WT and R164H, R164S variants, calculated from the equilibrium simulations.
